# Assessing seasonal demographic covariation to understand environmental-change impacts on a hibernating mammal

**DOI:** 10.1101/745620

**Authors:** Maria Paniw, Dylan Childs, Kenneth B Armitage, Daniel T Blumstein, Julien Martin, Madan K. Oli, Arpat Ozgul

## Abstract

Natural populations are exposed to seasonal variation in environmental factors that simultaneously affect several demographic rates (survival, development, reproduction). The resulting covariation in these rates determines population dynamics, but accounting for its numerous biotic and abiotic drivers is a significant challenge. Here, we use a factor-analytic approach to capture partially unobserved drivers of seasonal population dynamics. We use 40 years of individual-based demography from yellow-bellied marmots (*Marmota flaviventer*) to fit and project population models that account for seasonal demographic covariation using a latent variable. We show that this latent variable, by producing positive covariation among winter demographic rates, depicts a measure of environmental quality. Simultaneous, negative responses of winter survival and reproductive-status change to declining environmental quality result in a higher risk of population quasi-extinction, regardless of summer demography where recruitment takes place. We demonstrate how complex environmental processes can be summarized to understand population persistence in seasonal environments.

## INTRODUCTION

Effects of environmental change on survival, growth, and reproduction are typically investigated based on annual transitions among life-history stages in structured population models (Salguero-Gómez et al., 2016; Paniw et al., 2018). However, all natural ecosystems show some level of seasonal fluctuations in environmental conditions, and numerous species have evolved life cycles that are cued to such seasonality (Ruf et al., 2012; Varpe, 2017). For example, most temperate- and many arid-environment species show strong differences in survival and growth among seasons, with reproduction being confined mostly to one season (Childs et al., 2011; Rushing et al., 2017; Woodroffe et al., 2017). Species with highly adapted, seasonal life cycles are likely to be particularly vulnerable to environmental change, even if they are relatively long-lived (Jenouvrier et al., 2012; Campos et al., 2017; Paniw et al., 2019). This is because adverse environmental conditions in the non-reproductive season may carry-over and negate positive environmental effects in the reproductive season in which key life-history events occur (Marra et al., 2015). For instance, in species where individual traits such as body mass determine demographic rates, environment-driven changes in the trait distribution in one season can affect trait-dependent demographic rates in the next season (Bassar et al., 2016; Paniw et al., 2019). Investigating annual dynamics, averaged over multiple seasons, may, therefore, obscure the mechanisms that allow populations to persist under environmental change.

Despite the potential to gain a more mechanistic view of population dynamics, modeling the effects of seasonal environmental change is an analytically complex and data-hungry endeavor (Benton et al., 2006; Bassar et al., 2016). This is in part because multiple environmental factors that change throughout the year can interact with each other and individual-level (e.g., body mass) or population-level factors (e.g., density dependence) to influence season-specific demographic rates (Benton et al., 2006; Lawson et al., 2015; Ozgul et al., 2007; Paniw et al., 2019; Töpper et al., 2018). One major analytical challenge for ecologists is that typically only a small subset of the numerous biotic and abiotic drivers of important life-history processes are known and measured continuously (Teller et al., 2016); and this challenge is amplified in seasonal models where more detail on such drivers may be required while biological processes such as hibernation are cryptic to researchers (van de Pol et al., 2016). Assessing whether the available information provides meaningful measures of biological processes is another challenge. Nonlinear interactions among the myriad of biotic and abiotic factors are common in nature, and teasing apart their effects on natural populations requires detailed and long-term data (Benton et al., 2006; Paniw et al., 2019), which is not available for most systems (Salguero-Gómez et al., 2015; 2016).

Overcoming the challenges in parameterizing seasonal population models is important because a robust projections of such models require assessing the simultaneous effects of biotic and abiotic factors on several demographic rates, causing the latter to covary within and among seasons (Maldonado-Chaparro et al., 2018; Paniw et al., 2019). Positive environment-driven covariation in demographic rates can amplify the population-level effects of environmental change. For instance, Jongejans et al. (2010) demonstrated that positive covariation in survival and reproduction in several plant populations magnified the effect of environmental variability on population dynamics and increased extinction risk. On the other hand, antagonistic demographic responses, either due to intrinsic tradeoffs or opposing effects of biotic/abiotic factors, can buffer populations from environmental change (Knops et al., 2007; Van de Pol et al., 2010); for instance, when population-level effects of decreased reproduction are offset by increases in survival or growth (Connell & Ghedini, 2015; Reed et al., 2013; Villellas et al., 2015). Thus, explicit consideration of patterns in demographic covariation can allow for a fuller picture of population persistence in a changing world. Such a consideration remains scarce (Ehrlén & Morris, 2015; Ehrlén et al., 2016; but see Bassar et al., 2016; Compagnoni et al., 2016).

Here, we investigated the population-level effects of seasonal covariation among trait-mediated demographic rates (*i.e.*, collectively referred to as demographic processes), capitalizing on 40 years (1976-2016) of individual-based data from a population of yellow-bellied marmots (*Marmota flaviventer*). Our main aims were to (i) efficiently model demographic covariation in the absence of knowledge on its underlying drivers and (ii) characterize the seasonal mechanisms through which this covariation affects population viability. Yellow-bellied marmots have adapted to a highly seasonal environment; individuals spend approximately eight months in hibernation during the cold winter (September/October-April/May), and use the short summer season (April/May-September/October) to reproduce and replenish fat reserves (Fig. 1). One challenge that the marmot study shares with numerous other natural systems is the identification of key proximal biotic and abiotic factors driving population dynamics. In marmots such factors are numerous and affect population dynamics through complex, interactive pathways (Maldonado-Chaparro et al., 2017; Oli & Armitage, 2004), which include interactions with phenotypic-trait structure (Ozgul et al., 2010). As a result, measures of environmental covariates (e.g., temperature or resource availability) have previously shown little effect on the covariation of marmot demographic processes (Maldonado-Chaparro et al., 2018). To address this challenge, we used a novel method, a hierarchical factor analysis (Hindle et al., 2018), to model the covariation of demographic processes as a function of a shared latent variable, quantified in a Bayesian modeling framework. We then built seasonal stage-, mass-, and environment-specific integral projection models (IPMs; Ellner et al., 2016) for the marmot population, which allowed us to simultaneously project trait distributions and population dynamics across seasons. We used prospective stochastic perturbation analyses and population projections to assess how the observed demographic covariation mediated population viability.

**Figure 1:**
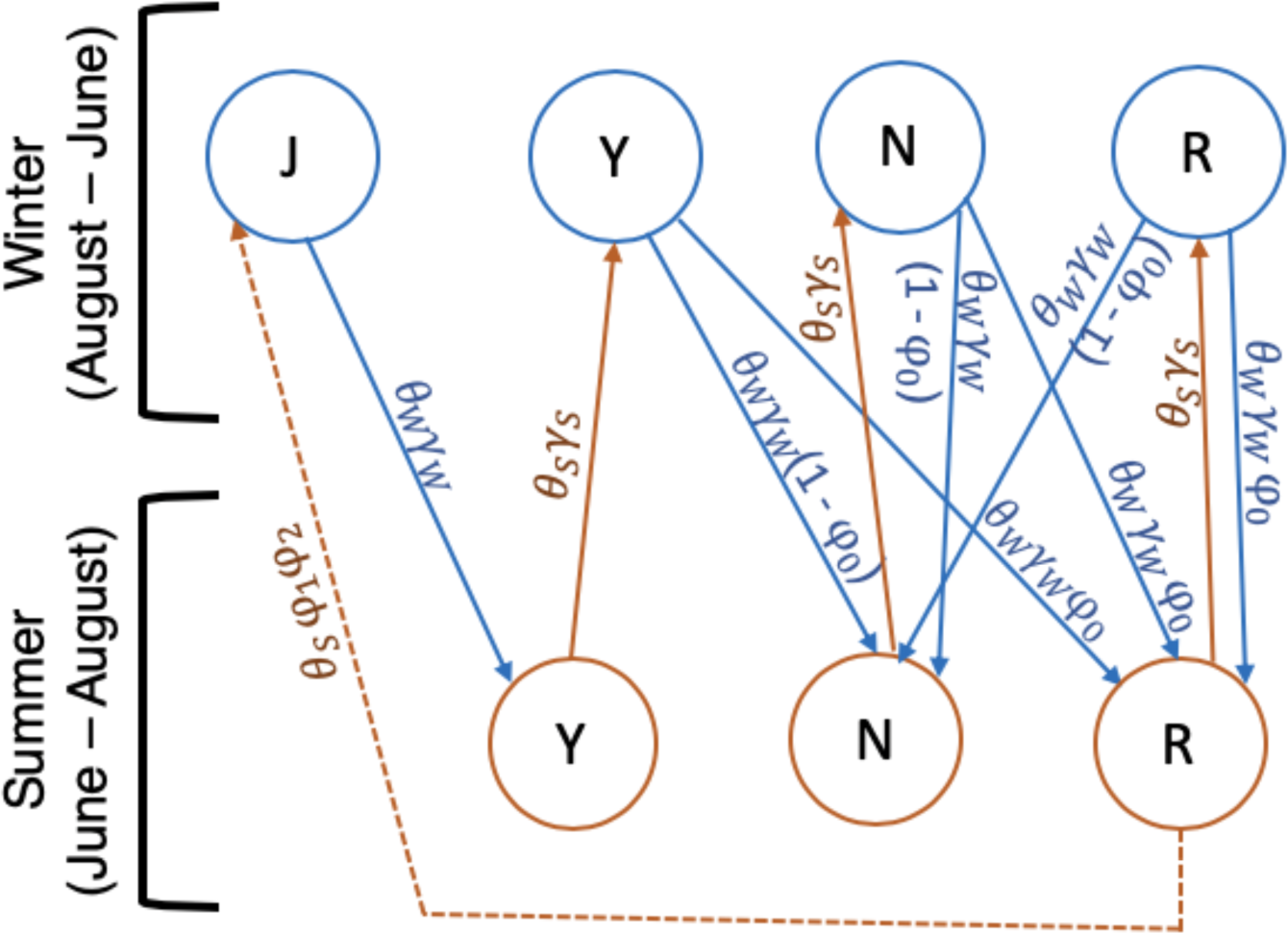
Seasonal life-cycle transitions modelled for yellow-bellied marmots. The two seasons correspond to the main periods of mass loss (winter) and gain (summer). Solid and dashed arrows represent discrete-time stage transitions and recruitment, respectively. Transitions among winter (W) and summer (S) stages (marked by arrows in different colors) depend on demographic rates (survival [*θ*], reproduction [*φ*_0_], and recruitment [*φ*_1_]) and trait transitions (mass change [*γ*], and offspring mass [*φ*_2_]). Stages are: juveniles, J, yearlings, Y, non-reproductive adults, N, and reproductive adults, R. All stage-specific demographic rates and trait transitions are modeled using generalized linear mixed effects models in a Bayesian framework and include body mass and a common latent variable representing environmental quality as covariates.

## METHODS

### Study species

Yellow-bellied marmots are an ideal study system to assess the effects of seasonal covariation in demographic rates on population viability. These large, diurnal, burrow-dwelling rodents experience strong seasonal fluctuations in environmental conditions, and their seasonal demography has been studied for > 40 years (Armitage, 2014). Our study was conducted in the Upper East River Valley near the Rocky Mountain Biological Laboratory, Gothic, Colorado (38° 57’ N, 106° 59’ W). Climatic conditions in both winter and summer have been shown to influence reproduction and survival in the subsequent season (Lenihan & Van Vuren, 1996; Van Vuren & Armitage, 1991). In addition, predation is major cause of death in the active summer season (Van Vuren, 2001; Maldonado-Chaparro et al., 2017) and may be particularly severe shortly before (Bryant & Page, 2005) or after hibernation (Armitage, 2014), especially in year with heavy snow (Blumstein, pers. obs.). The effects of these factors on the demography of yellow-bellied marmots are mediated through body mass, with heavier individuals more likely to survive hibernation, reproduce in summer, and escape predation (Armitage et al.,1976; Ozgul et al., 2010). Population dynamics of marmots are therefore likely to be susceptible to changes in seasonal patterns of biotic and abiotic drivers. However, numerous interacting climatic factors, such as temperature extremes and length of snow cover, determine both winter and summer environmental conditions. The effects on marmot demography of these climatic factors, and of interactions between climate and predation (the latter mostly a cryptic process) have been shown to be difficult to disentangle (Schwartz & Armitage, 2002; Schwartz & Armitage, 2005).

### Seasonal demographic rates and trait transitions

For this study, we focused on the population dynamics of eight major colonies continuously monitored since 1976 (Armitage, 2014; Supporting Material S1). Each year, marmots were live-trapped throughout the growing season in summer (and ear-tagged upon first capture), and their sex, age, mass, and reproductive status were recorded (Armitage & Downhower, 1974; Schwartz et al., 1998). All young males disperse from their natal colonies, and female immigration into existing colonies is extremely rare; as such, local demography can be accurately represented by the female segment of the population (Armitage, 2010). Thus, we focused on seasonal demographic processes of females only. We classified female marmots by age and reproductive status: juveniles (< 1 year old), yearlings (1 year old), and non-reproductive (≥ 2 years old; not observed pregnant or with offspring) and reproductive adults (≥ 2 years old; observed pregnant or with offspring) (Armitage & Downhower, 1974).

We determined demographic rates (survival, reproduction, and recruitment) for two discrete growing seasons: winter (August - June) and summer (June - August) (Fig. 1), delineating the main periods of mass loss and gain, respectively (Maldonado-Chaparro et al., 2017). We assumed that females that permanently disappeared from a colony had died. This measure of apparent survival may overestimate the death of yearlings in the summer, which disperse from their natal colonies (Van Vuren & Armitage, 1994). At the same time, the intensive trapping protocol ensured a high capture probability of yearlings (Oli & Armitage, 2004), decreasing the discrepancies between their apparent and true survival.

Female marmots give birth to one litter from mid-May to mid-June. In our population model, females ≥ 2 year of age that survived the winter were considered reproductive adults at the beginning of summer if they were observed to be pregnant or with pups, or non-reproductive adults otherwise (Fig. 1). Upon successful reproduction, weaned offspring emerge from burrows ca. 35 days after birth (Armitage et al., 1976); we therefore defined recruitment as the number of female juveniles weaned by reproductive females that survive the summer (Fig. 1). The sex ratio of female:male recruits was assumed to be 1:1 (Armitage & Downhower, 1974). Observations and pedigree analyses allowed us to determine the mother of each new juvenile recruited into the population (Ozgul et al., 2010).

To assess changes in body mass from one season to the next, we estimated body mass of every female at the beginning of each season: June 1 (beginning of the summer season when marmots begin foraging) and August 15 (beginning of the winter season in our models when foraging activity decreases). Mid-August is the latest that body mass for the vast majority of individuals can be measured and has been shown to be a good estimate of pre-hibernation mass (Maldonado-Chaparro et al., 2017). Body-mass estimates on the two specific dates were estimated using linear mixed effect models. These models were fitted for each age class and included the fixed effect of day-of-year on body mass, and the random effects of year, site and individual identity on the intercept and on the day-of-year slope (for details see Ozgul *et al*., 2010; Maldonado-Chaparro et al., 2017). Body mass of juvenile females was estimated for August 15.

### Modelling covariation in demographic processes – latent-variable approach

We jointly modeled all seasonal demographic and mass change rates (*i.e.*, demographic processes) as a function of stage and body mass - or mother’s mass in the case of juvenile mass - at the beginning of a season, using a Bayesian modeling framework (Table 1; Supporting Material S1). All mass estimates were cube-root transformed to stabilize the variance and improve the normality of the residuals in the Gaussian submodels (Maldonado-Chaparro et al., 2017). We fitted all demographic-process submodels as generalized linear mixed effects models (GLMMs). We assumed a binomial error distribution (logit link function) for the probability of winter (*θ*_W_) and summer (*θ*_S_) survival and of probability of reproducing (*i.e.*, being in the reproductive adult stage at the beginning of summer; φ_0_); a Poisson error distribution (log link function) for the number of recruits (φ_1_); and a Gaussian error distribution (identity link) for the masses (*z*\*) at the end of each season (Table 1). Mass-change (*i.e.*, mass gain or loss) rates (*γ*) were then defined as functions of current (*z*) and next (*z*\*) mass using a normal probability density function. For the juvenile mass distribution (φ_2_), the density function depended on the mother’s mass (*z*_*M*_) (see below; Supporting Material S2).

**Table 1:** Parameterization of the most parsimonious models describing winter (W) and summer (S) demographic processes in marmots. The distributions B, N, and P correspond to the Bernoulli, normal, and Poisson distributions, respectively. *Stage* – life cycle stage. *Q* – latent environmental variable. *z* – season-specific mass. *z*_*M*_ –mass of the mother.

| Demographic process | Function | Likelihood distribution |
| --- | --- | --- |
| **Winter (W):** |  |  |
| Survival ($\theta_W$) | $\text{logit}(\theta_W) = \alpha_{0\theta W} + \alpha_{a\theta W}[\text{stage}] + \beta_{z\theta W} \times z + \beta_{q\theta W} \times Q_y[\text{year}] + \varepsilon_{y\theta W}[\text{year}]$ | $B(\theta_W)$ |
| Mass next ($z_W^*$) | $z_W^* = \alpha_{0z^*W} + \alpha_{az^*W}[\text{stage}] + (\beta_{zz^*W} + \beta_{zaz^*W}[\text{stage}]) \times z + \beta_{qz^*W} \times Q_y[\text{year}] + \varepsilon_{yz^*W}[\text{year}]$ | $\mathcal{N}(z_W^*, \tau_{z^*W})$ |
| Reproduction ($\varphi_0$) | $\text{logit}(\varphi_0) = \alpha_{0\varphi_0} + \alpha_{a\varphi_0}[\text{stage}] + \beta_{z\varphi_0} \times z + \beta_{q\varphi_0} \times Q_y[\text{year}] + \varepsilon_{y\varphi_0}[\text{year}]$ | $B(\varphi_0)$ |
| **Summer (S):** |  |  |
| Survival ($\theta_S$) | $\text{logit}(\theta_S) = \alpha_{0\theta S} + \alpha_{a\theta S}[\text{stage}] + \beta_{z\theta S} \times z + \beta_{q\theta S} \times Q_y[\text{year}] + \varepsilon_{y\theta S}[\text{year}]$ | $B(\theta_S)$ |
| Mass next ($z_S^*$) | $z_S^* = \alpha_{0z^*S} + \alpha_{az^*S}[\text{stage}] + (\beta_{zz^*S} + \beta_{zaz^*S}[\text{stage}]) \times z + \beta_{qz^*S} \times Q_y[\text{year}] + \varepsilon_{yz^*S}[\text{year}]$ | $\mathcal{N}(z_S^*, \tau_{z^*S})$ |
| Number of recruits ($\varphi_1$) | $\log(\varphi_1) = \alpha_{0\varphi_1} + \beta_{z\varphi_1} \times z + \beta_{q\varphi_1} \times Q_y[\text{year}] + \varepsilon_{y\varphi_1}[\text{year}]$ | $P(\varphi_1)$ |
| Juvenile mass ($z_J^*$) | $z_J^* = \alpha_{0z^*J} + \beta_{zz^*J} \times z_M + \beta_{qz^*J} \times Q_y[\text{year}] + \varepsilon_{yz^*J}[\text{year}]$ | $\mathcal{N}(z_J^*, \tau_{z^*J})$ |

To model temporal covariation in seasonal demography in the absence of explicit knowledge on key biotic or abiotic drivers of this covariation, we used a factor-analytic approach. This approach has recently been proposed by Hindle and coauthors (2018) as a structured alternative to fit and project unstructured covariances among demographic processes when factors explaining these covariances are not modeled. We implemented this novel approach parameterizing a model-wide latent variable (*Q*_*y*_) which affected all demographic processes in a given year (*y*) (for details see Supporting Material S1 and Hindle *et al*., 2018). *Q*_*y*_ was incorporated as a covariate in all seven demographic-process submodels (Table 1). Year-specific values of *Q*_*y*_ were drawn from a normal distribution with mean = 0 and SD =1. The associated *β*_*q*_ slope parameters then determine the magnitude and sign of the effect of *Q*_*y*_ on a given, season-specific demographic process (Table 1). To make the Bayesian model identifiable, we constrained the standard deviation of *Q*_*y*_ to equal 1 and arbitrarily set the *β*_*q*_ for summer survival (*θ*_S_) to be positive. The *β*_*q*_ of the remaining submodels can, therefore, be interpreted as correlations of demographic processes with *θ*_S_.

Aside from the latent variable *Q*_*y*_ simultaneously affecting all demographic processes, we included a random year effect (*ε*_*Ysubmodel*_) as a covariate in each submodel. While *Q*_*y*_ captured demographic covariation, the year effect accounted for additional temporal variation of each demographic process not captured by *Q*_*y*_. We also tested for the effect of population density (measured as total abundance, abundance of adults, or abundance of yearling and adults) in all submodels. However, like previous studies, we could not detect any clear density effects (Armitage, 1984; Maldonado-Chaparro et al., 2018).

The prior distributions of the Bayesian model and posterior parameter samples obtained are detailed in Supporting Material S1. For each demographic-process submodel, we chose the most parsimonious model structure by fitting a full model that included all covariates (mass, stage, and *Q*_*y*_) and two-way interactions between mass and stage and stage and *Q*_*y*_, and retaining only those parameters for which the posterior distribution (± 95 % C.I.) did not overlap 0 (Table 1; Table S1.1).

### Interpreting demographic covariation: latent variable as a measure of environmental quality

The latent variable, *Q*_*y*_, effectively captured the covariation among the demographic processes (Supporting Material S1); therefore, using one latent variable across both seasons was sufficient. Our GLMMs showed a strong effect of *Q*_*y*_ on winter but not summer demographic processes. This effect was positive for all winter demographic processes, as evidenced by the positive *β*_*q*_ (Table S1.1). The *β*_*q*_ for demographic processes in the summer, however, were comparatively small and were not significantly different from 0 (95 % posterior C.I.s overlapped 0). The positive *β*_*q*_ indicate that *Q*_*y*_ effectively estimates the overall annual environmental quality or suitability, capturing both biotic and abiotic processes. A positive value of *Q*_*y*_ then depicts an environmental condition at a given time point that increases winter survival and probability of reproducing and decreases mass loss (Hindle et al., 2018). The variation in *Q*_*y*_ was in part explained by environmental variables measured at the study site, but was unrelated to population density (Supporting Material S1). Negative values of *Q*_*y*_ were associated with longer and more severe winters and a higher snow cover, while positive *Q*_*y*_ indicated warmer winters and springs. However, as the environmental variables explained < 50 % of the variation in *Q*_*y*,_ the latent variable captures multivariate, partly unobserved biotic and abiotic processes into a simple, univariate measure of how bad (*Q*_*y*_ < 0) or good (*Q*_*y*_ > 0) environmental conditions are likely to affect marmot demography.

Aside from the effects of environmental quality, our models are consistent with previous findings on the importance of body mass and stage on yellow-bellied marmot demography (Maldonado-Chaparro et al., 2017; Ozgul et al., 2010). The most parsimonious GLMMs (Table S1.1) showed a positive effect of mass on all demographic processes, with the weakest effect of mass on summer survival (*θ*_S_) of reproductive adults. Survival, in particular *θ*_S_, was highest for reproductive adults; reproduction was also highest for adults that reproduced before (Fig. S1.5).

### Seasonal Integral Projection Models

We used the most parsimonious models of demographic processes (Table 1) to parameterize density-independent, stage-mass-structured, seasonal and environment-specific Integral Projection Models (IPMs) (Easterling et al., 2000; Ellner et al., 2016). For each stage *a*, the IPMs track the number of individuals (*n*_*a*_) in the mass range [*z, z*+d*z*] at time *t*. The fate of these individuals at time *t*+1 is described by a set of coupled integral equations, which differ for each season and are a function of the latent environmental variable *Q*_*y*_. In the winter season, individuals can survive (*θ*_w_) and change mass (*γ*_W_) according to their stage, mass, and environment. Conditional on survival, juveniles (J) transition to yearlings (Y), while all other stages are distributed to either the reproductive (R) or non-reproductive (N) adult stage at the beginning of summer, depending on the stage-specific probability of reproducing (φ_0_). During the summer season, individuals in stages Y, N, and R survive (*θ*_S_) and change mass (*γ*_S_) according to their stage and mass at the beginning of summer and according to the environment; but, in summer, transitions to another stage do not occur. Reproductive individuals (R) of a given mass also produce φ_1_/2 female juveniles (J), *i.e.*, half of the total number of recruits. Female recruits are distributed across *z* mass classes by the end of summer, given by φ_2_. The mathematical descriptions of the IPMs for the winter and summer seasons are provided in Supporting Material S2. Our population model assumes that past conditions affecting individuals are captured by the current mass distribution and are propagated through time, allowing us to assess trait- and stage-mediated demographic processes (Ozgul et al., 2010).

We numerically integrated the summer and winter IPMs using the ‘midpoint rule’ (Easterling et al., 2000) with upper and lower integration limits of 7.8 (472 g) and 17.1 (5000 g), respectively. To avoid unintended eviction of individuals from the model (*i.e.*, for a given mass class *z*, the sum of the probabilities to transition to *z\** < 1), we applied a constant correction (*i.e.*, equally redistributing evicted individuals among all *z\**) when constructing the IPMs as suggested in Merow *et al.*, (2014) (see also Williams et al., 2012). For each stage-specific IPM, we chose a bin size of 100 (*i.e*., dividing masses into 100 classes), as further increasing the bin size did not significantly improve the precision of estimates of the long-term population growth rate. The IPMs we constructed accurately reproduced observed population dynamics from 1976-2016 (Supporting Material S2).

### Sensitivity of population dynamics to seasonal demographic processes: prospective perturbations

Changes in population dynamics in response to changes in environmental fluctuations are determined by the response of demographic processes to the environment and, in turn, of population dynamics to demographic processes (Maldonado-Chaparro et al., 2018). To explore these two sources of variation in the long-term fitness of the marmot population, we first quantified the proportional change in the demographic processes (Table 1) to changes in *Q*_y_, *i.e.*, ∂(log ρ)/∂*Q*_*y*_, where ρ is a demographic process. We calculated these elasticities for different values of *Q*_y_ (from −1 to 1), increasing each value by 0.01 and keeping mass at its stage-specific average and *ε*_*Y*_ fixed to 0. To assess the effect of parameter uncertainty on our estimates, we repeated these calculations for a sample of 1000 parameter values drawn from the posterior distribution (Paniw et al., 2017).

We next assessed which demographic processes most affected the stochastic population fitness under observed (1976-2016) environmental fluctuations. We used a simulation of 100,000 years to assess the stochastic population growth rate, log *λs*, a measure of fitness (see Supporting Material S3 for details; see section below for short-term viability simulations). During the simulation, we calculated the elasticity of log *λs* to changes in the 40-year observed mean 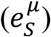 and standard deviation 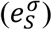 of stage-specific demographic processes; we adapted the approach described in Ellner *et al*. (2016; chapter 7) to evaluate the relative effects of these changes on log *λs* (see S3 for details). The two elasticities quantify the strength of selection pressures on lower-level vital rates in stochastic environments (Haridas & Tuljapurkar, 2005; Rees & Ellner, 2009). We repeated the elasticity calculations for a sample of 100 parameter values from the posterior distribution.

### Population viability under changes in environmental quality

To assess how the combined effects of (i) seasonal demographic responses to environmental fluctuations and (ii) population sensitivity to seasonal demography impact population viability, we simulated population dynamics under environmental change. We ran 200 independent simulations each projecting population dynamics for 50 years. The projections were based on several scenarios of changes in the distribution of environmental quality, *Q*_*y*_, corresponding to changes in the average and standard deviation of winter length and harshness as well as unobserved environmental drivers. We first created base simulations (*i.e.*, no environmental change) where *Q*_*y*_ was picked from a normal distribution with *μ*_*Q*_ = 0 and σ_*Q*_ = 1 across all demographic processes. This was appropriate, as we found no indication of temporal autocorrelation in *Q*_*y*_ (Supporting Material S1). Next, we approximated random future fluctuations in *Q*_*y*_ under different average environmental conditions. To do so, we sampled *Q*_*y*_ from a normal distribution fixing the average environmental quality (*μ*_*Q*_ = −1, −0.5, 0.5, 1) and its variation (σ_*Q*_ = 0.6, 1.2) over the 50 years of projections. We then explored how a trend in *μ*_*Q*_ would affect viability and mass distribution. To do so, we decreased the four *μ*_*Q*_ by 0.01 in each year of the projections, keeping σ_*Q*_ unaltered. We also explored population-level effects of future increases in the temporal autocorrelation in *Q*_*y*_ as detailed in Supporting Material S4. All simulations were repeated for a random sample of 1,000 parameters from the posterior distribution to account for parameter uncertainty.

For all environmental-change scenarios, we recorded the probability of quasi-extinction across the 200 simulations. Quasi-extinction was defined conservatively as the number of non-juvenile individuals (*i.e.*, yearlings and non-reproductive and reproductive individuals) in the population to be < 4, which corresponded to 10 % of their lowest observed number.

## RESULTS

### Sensitivity of population dynamics to seasonal demographic processes

In accordance with the posterior distribution of *β*_*q*_ parameters, which did not cross 0 for winter demographic processes, only winter demographic processes were significantly affected by small changes in *Q*_y_ (Fig. 2). Among the winter demographic processes, changes in *Q*_y_ affected reproduction across stages the most, followed by survival of juveniles (Fig. 2).

**Figure 2:**
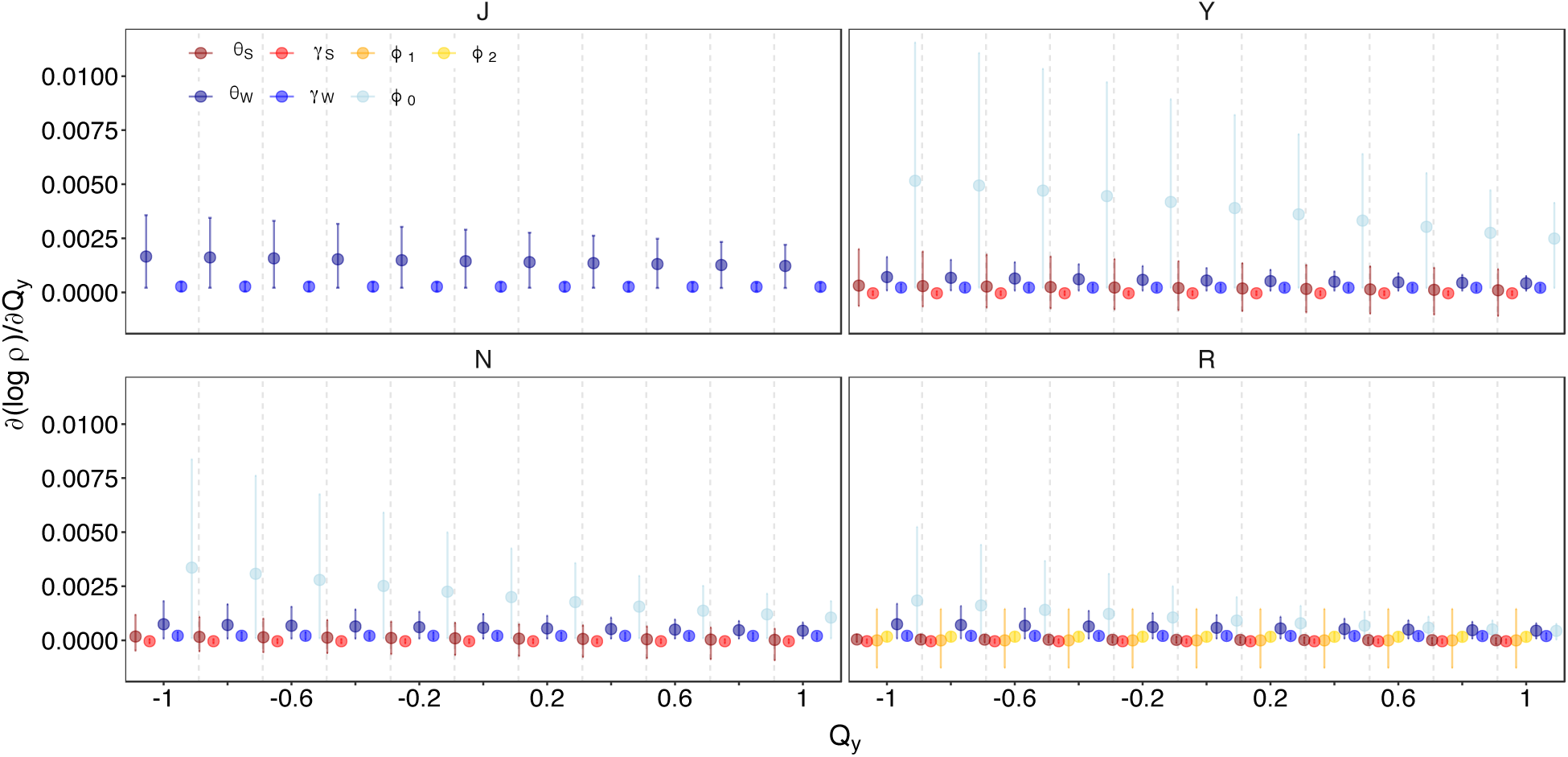
The sensitivity of seasonal demographic processes to environmental quality in marmots. Sensitivity is assessed as proportional changes in demographic processes, ρ, as environmental quality, *Q*_*y*_, increases slightly. This sensitivity is measured with respect to different average values of *Q*_y_ and across four different life-cycle stages: juveniles (J), yearlings (Y), non-reproductive adults (N), and reproductive adults (R). The demographic processes include winter (W; blue color tones) and summer (S; red color tones) survival (*θ*) and mass change (*γ*); and probability of reproducing (φ_0_), recruitment (φ_1_), and juvenile mass (φ_2_). Points and error bars show averages ± 95 % C.I. across 1,000 posterior parameter samples obtained from the Bayesian population model.

While environmental quality affected winter demographic processes only, our prospective perturbation analyses showed that winter and summer demography equally determine long-term population fitness. Stochastic elasticity analyses (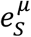and 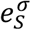) showed that relative increases in the mean (µ) of winter (*θ*_W_) and summer (*θ*_S_) survival for reproductive adults (R), would lead to substantial relative increases of the stochastic population growth rate, log*λs* (Fig. 3a). Highest, positive 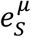 were found at intermediate and large mass classes, and 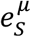 was negative for small masses when mass changes (*γ*) and offspring mass (φ_2_) were perturbed (Fig. S3.1a in Supporting Material S3). This explained the overall small 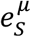 for *γ* and φ_2_ summed over all mass classes (Fig. 3a). Overall, relative changes in log*λs* due to increases in the standard deviation of demographic processes 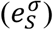 were much smaller compared to 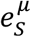 (Fig. 3b) and didn’t differ significantly between vital rates, as 95 % posterior C.I. crossed 0 (Fig. S3.1b).

**Figure 3.**
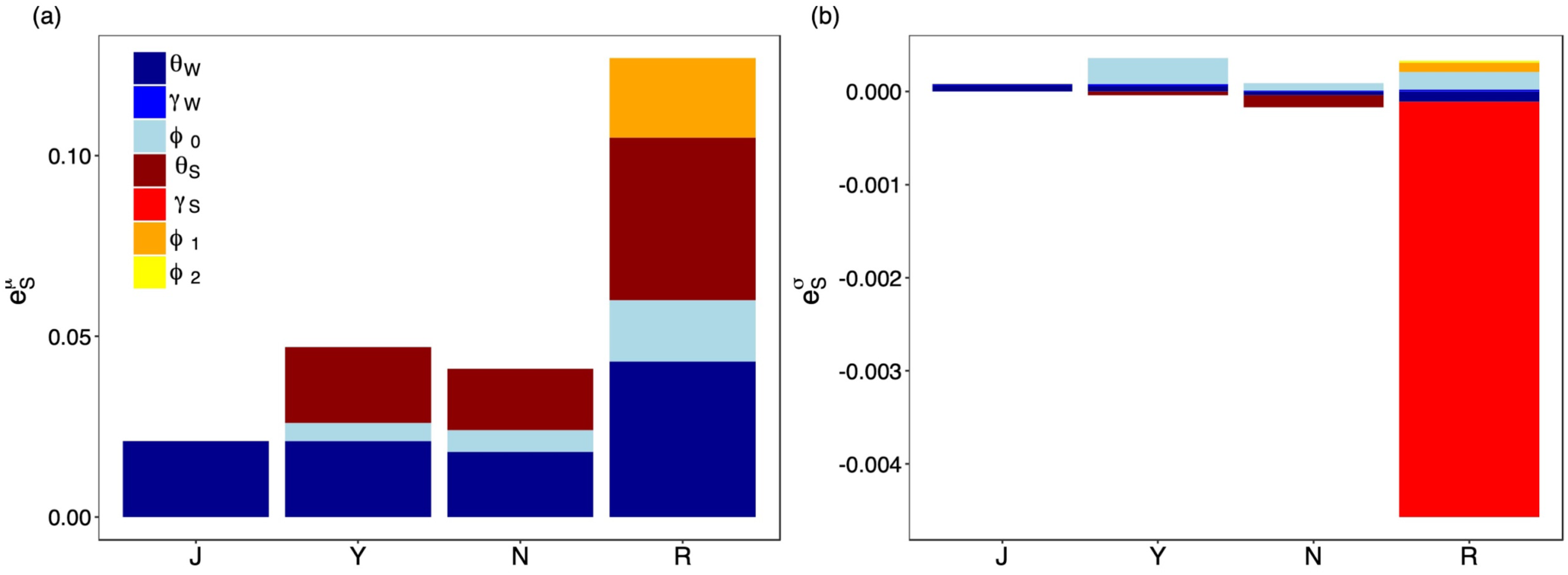
Sensitivity of the average long-term population fitness to changes in the average and variability of demographic processes modeled for the yellow-bellied marmots. The sensitivity measure is obtained analytically as elasticities (e) of the stochastic population growth rate, log λs, to changes in (a) the mean (*μ*) and (b) standard deviation (σ) of stage-specific demographic processes summed over all mass classes. Stages are juveniles (J), yearlings (Y), non-reproductive adults (N), and reproductive adults (R). Demographic processes include winter (W) and summer (S) survival (*θ*) and mass change (*γ*); reproduction (φ_0_); recruitment (φ_1_), and offspring mass distribution (φ_2_). Elasticities were calculated at the mean posterior values of parameters obtained from the Bayesian demographic model.

### Population viability under changes in environmental quality

While population fitness was equally sensitive to demographic processes over winter and summer, environmental fluctuations strongly affected viability through winter demography. Using base simulations (*i.e.*, obtaining *Q*_*y*_ from a normal distribution with *μ*_*Q*_ = 0 and σ_*Q*_ = 1), the probability of quasi-extinction, at an average of 0.1 [0.0, 0.3 C.I.] across posterior parameters, were relatively low. Simulations of population dynamics based on scenarios of environmental change, corresponding in part to changes in winter length and harshness, resulted in substantial changes to viability. Quasi-extinction decreased (0 at *μ*_*Q*_ = 1) and increased (0.9 [0.6, 1.0 C.I.] at *μ*_*Q*_ = −1), compared to base simulations, when the population experienced a high and low average environmental quality (*Q*_*y*_), respectively (Fig. 4). Average quasi-extinction further increased and its uncertainty across posterior parameters decreased when a declining trend in *Q*_*y*_ was simulated (Fig. S4.1). Changes in the standard deviation (Fig. 4) and autocorrelation (Fig. S4.2) of *Q*_*y*_ had comparatively little effect on quasi-extinction.

**Figure 4:**
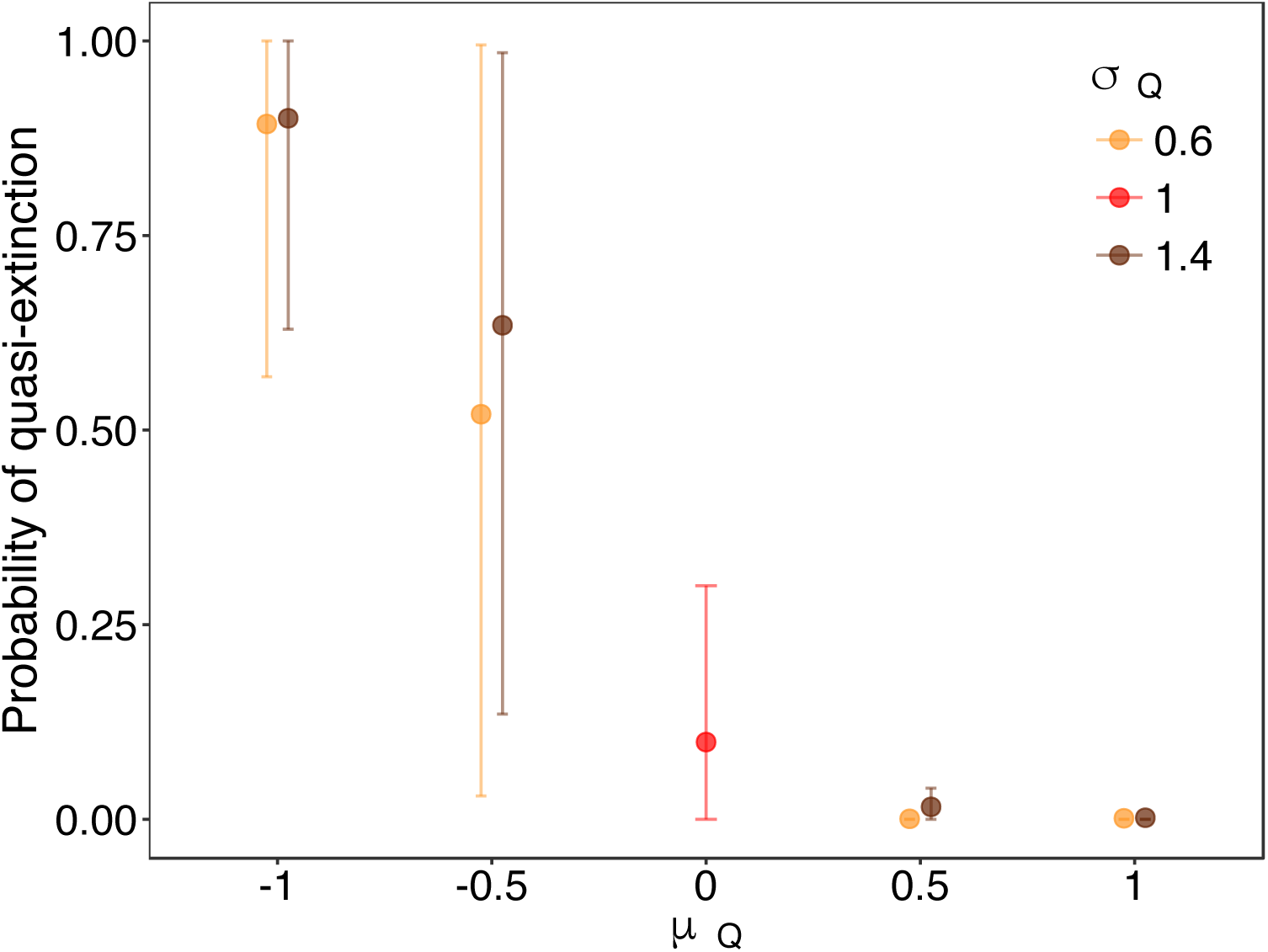
Probability of quasi-extinction (*i.e.*, < 4 non-juveniles in the population) of yellow-bellied marmots under different scenarios of environmental change. The scenarios consisted of projecting population dynamics for 50 years fixing a different mean (*μ*) and standard deviation (σ) of environmental quality (*Q*) in all demographic processes. Points and error bars show averages ± 95 % C.I. across 1,000 posterior parameter samples obtained from the Bayesian population model. Base simulations (*μ*_*Q*_ = 0; σ_*Q*_ = 1) are depicted in red.

## DISCUSSION

One important pathway through which environmental change can act on population dynamics is through seasonal direct and carry-over effects on survival, development, and reproduction (Harrison et al., 2010; Paniw et al., 2019). These effects, however, are often cryptic and therefore difficult to quantify in ecological models (Hindle et al., 2019). We use a novel, factor-analytic approach to efficiently quantify partially unobserved environment-demography relationships. This approach allows us to investigate how positive responses in several demographic processes to winter environmental conditions can drive annual population dynamics in a winter-adapted mammal. The sensitivity to winter conditions occurs despite the fact that offspring are recruited in summer and both summer and winter demographic processes determine population fitness. As whole-year, population-level effects of environmental change can be filtered by season-specific processes in the absence of density-dependent feedbacks, we highlight that the assessment of such processes allows for a mechanistic understanding of population persistence (Picó et al., 2002; Paniw et al., 2019).

In marmots, as in numerous other populations (Bassar et al., 2016; Jenouvrier et al., 2018), seasonal demographic processes play an important role in life-cycle dynamics (Armitage, 2017). Our prospective perturbations show that changes in both mean winter and summer survival of reproductive adults have the strongest effect on population fitness, confirming the critical role of this life-cycle stage (Ozgul et al., 2009). At the same time, environmental conditions do not affect adult survival or other demographic processes in the same way throughout the year. That is, although the environment has been shown to drive particularly recruitment in numerous temperate species (e.g., Bonardi et al., 2017; Nouvellet et al., 2013), such effects are not evident in marmots; here, a higher annual environmental quality, which increases all winter demographic processes, shows little impact on summer demography, including recruitment. In turn, only these joint responses of winter demographic processes to environmental quality determine population persistence under environmental change.

The complex, partially unmeasured environmental processes that cause positive covariation in seasonal demographic processes can be effectively captured using a univariate, latent measure of environmental quality. In our study, this latent quality correlated better with observed annual population growth than any measured environmental variable (Supporting Material S1). In part, a good quality depicts shorter and milder winters. Milder winters increase food availability and the time available for vigilance, thereby decreasing predation risk (Van Vuren, 2001), especially just before or after hibernation (*i.e.*, within our winter season) when such risk is severe (Armitage, 2014). Predation risk in early spring also increases under high snow cover, as marmots, including more experienced adult females, cannot easily retreat to their burrows (Blumstein, pers. obs.). Predation is however cryptic in the system (Van Vuren, 2001). Capturing the effects of unobserved environmental variation, including predation, the latent-variable approach appears to be a promising alternative to modeling seasonal demographic processes under limited knowledge of their drivers (Evans & Holsinger, 2012; Hindle et al., 2019; Hindle et al., 2018). We note that this approach may find limited applications in cases where environment-demography relationships are more complex than in the yellow-bellied marmots and include negative demographic covariation (e.g., due to opposing environmental effects on demographic rates or tradeoffs between these rates). However, positive covariation in demographic patterns is common (Jongejans et al., 2010; Paniw et al., 2019); and, given the short time series of most demographic datasets (Salguero-Gómez et al., 2015; 2016) or little knowledge on the actual environmental drivers of population dynamics (van de Pol et al., 2016; Teller et al., 2016), the factor-analytic approach can be particularly useful in comparative studies.

The seasonal effects of environmental quality on population persistence must be understood in terms of the role of reproductive females in the marmot population (Ozgul et al., 2009). In our simulations, shorter and less sever winters (*i.e*., a good winter quality), would result in more reproductive females in the summer (Armitage et al., 2003). In turn, summer survival and recruitment by these females are important to long- and short-term demography (Ozgul et al., 2009; Maldonado-Chaparro et al., 2018), but are not driven by environmental conditions. That is, although predation affects individuals in summer (Van Vuren, 2001), its effects are strongest on juveniles and yearlings, while adult females are little affected (Ozgul et al., 2006). At the same time, as is the case in other socially complex mammals (Morris et al., 2011), reproduction in yellow-bellied marmots is governed primarily by social interactions, in particular the behavior of dominant adult females (Armitage, 2010; Blumstein & Armitage, 1998). Even under optimal summer conditions, the reproductive output of the population may not increase as dominant females suppress reproduction in younger subordinates and therefore regulate the size of colonies (Armitage, 1991). Dominant females, in addition, may skip reproduction themselves if they enter hibernation with a relatively low mass (Armitage, 2017). Thus, the necessity of meeting the physiological requirements of hibernation profoundly affects life-history traits of yellow-bellied marmots that are expressed during the active season.

Unlike the effects of seasonal survival and reproduction, trait transitions between seasons had a smaller effect on annual population dynamics, even if winter mass changes were mediated by environmental quality. These relatively small effects are likely due to the fact that marmots compensate for winter mass loss with increased growth in the summer, creating a zero-net effect on annual trait change (Maldonado-Chaparro et al., 2017; 2018). Although the strength of compensatory effects may differ within seasons or among life-history stages (Monclús et al., 2014), such effects are common in rodents and other species that have a short window for mass gain (Morgan & Metcalfe, 2001; Orizaola et al., 2014), and highlight how assessing seasonal dynamics can provide a mechanistic understanding of population-level global-change effects (Bassar et al., 2016).

Under environmental change, the persistence of marmots was mostly affected by changes in mean environmental quality, whereas changes in the variance and temporal autocorrelation of the mean showed little effects. This supports previous conclusions that yellow-bellied marmots are partly buffered against increases in environmental variation (Maldonado-Chaparro et al., 2018; Morris et al., 2008) or autocorrelation (Engen et al., 2013). Further support for demographic buffering comes from the fact that changes in the mean environmental quality most strongly affected those demographic processes to which the stochastic population growth rate was least sensitive, *i.e*., yearlings gaining reproductive status. It is well known that in species where vital rates of adults are relatively buffered, juveniles are much more sensitive to environmental variation (Gaillard & Yoccoz, 2003; Jenouvrier et al., 2018). Our results indicate that demographic buffering (Pfister, 1998; Morris et al., 2008) likely persists across the seasonal environments and different masses for a high-altitude specialist.

Our results emphasize that declines in environmental quality in the non-reproductive season alone can strongly affect annual population dynamics of a mammal highly adapted to seasonal environments. Therefore, positive demographic covariation under environmental change may threaten populations even if it affects demographic process to which the stochastic growth rate is least sensitive, *i.e.*, processes that are under low selection pressure (Coulson et al., 2005; Iles et al., 2019). Studies that focus on the effects of environmental factors on single demographic processes that strongly affect both short- and long-term population dynamics may therefore underestimate the important role of seasonal demographic covariation.

Most species inhabit seasonal environments. Under global environmental change, it may therefore be critical to understand how seasonal patterns mediate persistence of natural populations. Novel methods such as the factor analytic approach allow researchers to overcome some challenges associated with more mechanistic approaches assessing population responses to environmental change, and we encourage more seasonal demographic analyses across different taxa.

## ACKNOWLEDGMENTS

We thank the many volunteers and researchers of Rocky Mountain Biological Laboratory for collecting and providing us with the data on individual life histories of yellow-bellied marmots. M.P. was supported by an ERC Starting Grant (33785) and a Swiss National Science Foundation Grant (31003A_182286) to A.O.; and by a Spanish Ministry of Economy and Competitiveness Juan de la Cierva-Formación grant FJCI-2017-32893; D.T.B. was supported by the National Geographic Society (grant # 8140-06), the UCLA Faculty Senate and Division of Life Sciences, a RMBL research fellowship, the National Science Foundation (IDBR-0754247, DEB-1119660 and 1557130 to D.T.B.; DBI 0242960, 0731346, and 1226713 to the RMBL).

## REFERENCES

Armitage, K. B. (1984). Recruitment in yellow-bellied marmot populations: kinship, philopatry, and individual variability. In J. O. Murie & G. R. Michener (Eds.), The Biology of Ground-Dwelling Squirrels (pp. 377–403), Lincoln: Univ. Nebraska Press.

Armitage, K. B. (1991). Social and population dynamics of yellow-bellied marmots: results from long-term research. Annual Review of Ecology and Systematics, 22, 379–407.

Armitage, K. B. (2010). Individual fitness, social behavior, and population dynamics of yellow-bellied marmots. I. Billick & M.V. Price (Eds.) The Ecology of Place: Contributions of Place-Based Research to Ecological Understanding (pp. 132–154), Chicago: University of Chicago Press.

Armitage, K. B. (2014). Marmot Biology: Sociality, Individual Fitness, and Population Dynamics. Cmbridge: Cambridge University Press.

Armitage, K. B. (2017). Hibernation as a major determinant of life-history traits in marmots. Journal of Mammalogy, 98, 321–331.

Armitage, K. B., Blumstein, D. T., & Woods, B. C. (2003). Energetics of hibernating yellow-bellied marmots (*Marmota flaviventris*). Comparative Biochemistry and Physiology. Part A, Molecular & Integrative Physiology, 134, 101–114.

Armitage, K. B., & Downhower, J. F. (1974). Demography of yellow-bellied marmot populations. Ecology, 55, 1233–1245.

Armitage, K. B., Downhower, J. F., & Svendsen, G. E. (1976). Seasonal changes in weights of marmots. The American Midland Naturalist, 96, 36–51.

Bassar, R. D., Letcher, B. H., Nislow, K. H., & Whiteley, A. R. (2016). Changes in seasonal climate outpace compensatory density-dependence in eastern brook trout. Global Change Biology, 22, 577–593.

Benton, T. G., Plaistow, S. J., & Coulson, T. N. (2006). Complex population dynamics and complex causation: devils, details and demography. Proceedings. Biological Sciences, 273, 1173–1181.

Blumstein, D. T., & Armitage, K. B. (1998). Life history consequences of social complexity a comparative study of ground-dwelling sciurids. Behavioral Ecology, 9, 8–19.

Bonardi, A., Corlatti, L., Bragalanti, N., & Pedrotti, L. (2017). The role of weather and density dependence on population dynamics of Alpine-dwelling red deer. Integrative Zoology, 12, 61–76.

Bryant, A. A. & Page, R. W. (2005). Timing and causes of mortality in the endangered Vancouver Island marmot (*Marmota vancouverensis*). Canadian Journal of Zoology, 83, 674–682.

Campos, F. A., Morris, W. F., Alberts, S. C., Altmann, J., Brockman, D. K., Cords, M., Pusey, A., Stoinski, T. S., Strier, K. B., & Fedigan, L. M. (2017). Does climate variability influence the demography of wild primates? Evidence from long-term life-history data in seven species. Global Change Biology, 23, 4907–4921.

Childs, D. Z., Coulson, T. N., Pemberton, J. M., Clutton-Brock, T. H., & Rees, M. (2011). Predicting trait values and measuring selection in complex life histories: reproductive allocation decisions in Soay sheep. Ecology Letters, 14, 985–992.

Compagnoni, A., Bibian, A. J., Ochocki, B. M., Rogers, H. S., Schultz, E. L., Sneck, M. E., Elderd, B. D., Iler, A. M., Inouye, D. W., Jacquemin, H., & Miller, T. E. X. (2016). The effect of demographic correlations on the stochastic population dynamics of perennial plants. Ecological Monographs, 86, 480–494.

Connell, S. D., & Ghedini, G. (2015). Resisting regime-shifts: the stabilising effect of compensatory processes. Trends in Ecology & Evolution, 30, 513–515.

Coulson, T., Gaillard, J.-M., & Festa-Bianchet, M. (2005). Decomposing the variation in population growth into contributions from multiple demographic rates. The Journal of Animal Ecology, 74, 789–801.

Easterling, M. R., Ellner, S. P., & Dixon, P. M. (2000). Size-specific sensitivity: applying a new structured population model. Ecology, 81, 694–708.

Ehrlén, J., & Morris, W. F. (2015). Predicting changes in the distribution and abundance of species under environmental change. Ecology Letters, 18, 303–314.

Ehrlén, J., Morris, W. F., von Euler, T., & Dahlgren, J. P. (2016). Advancing environmentally explicit structured population models of plants. The Journal of Ecology, 104, 292–305.

Ellner, S. P., Childs, D. Z., & Rees, M. (2016). Data-driven Modelling of Structured Populations: A Practical Guide to the Integral Projection Model. Springer.

Engen, S., Sæther, B.-E., Armitage, K. B., Blumstein, D. T., Clutton-Brock, T. H., Dobson, F. S., Festa-Bianchet, M., Oli, M. K., & Ozgul, A. (2013). Estimating the effect of temporally autocorrelated environments on the demography of density-independent age-structured populations. Methods in Ecology and Evolution, 4, 573–584.

Evans, M. E. K., & Holsinger, K. E. (2012). Estimating covariation between vital rates: a simulation study of connected vs. separate generalized linear mixed models (GLMMs). Theoretical Population Biology, 82, 299–306.

Gaillard, J.-M., & Yoccoz, N. G. (2003). Temporal variation in survival of mammals: A case of environmental canalization? Ecology, 84, 3294–3306.

Gamelon, M., Grøtan, V., Nilsson, A. L. K., Engen, S., Hurrell, J. W., Jerstad, K., Phillips, A. S., Røstad, O. W., Slagsvold, T., Walseng, B., Steneth, N. C., & Sæther, B.-E. (2017). Interactions between demography and environmental effects are important determinants of population dynamics. Science Advances, 3, e1602298.

Haridas, C. V., & Tuljapurkar, S. (2005). Elasticities in variable environments: properties and implications. The American Naturalist, 166, 481–495.

Harrison, X. A., Blount, J. D., Inger, R., Norris, R. D., & Bearhop, S. (2010). Carry-over effects as drivers of fitness differences in animals. The Journal of Animal Ecology, 80, 4–18.

Hindle, B. J., Pilkington, J. G., Pemberton, J. M., & Childs, D. Z. (2019). Cumulative weather effects can impact across the whole life cycle. Global Change Biology, in press.

Hindle, B. J., Rees, M., Sheppard, A. W., Quintana-Ascencio, P. F., Menges, E. S., & Childs, D. Z. (2018). Exploring population responses to environmental change when there is never enough data: a factor analytic approach. Methods in Ecology and Evolution, 147, 115.

Iles, D. T., Rockwell, R. F., & Koons, D. N. (2019). Shifting vital rate correlations alter predicted population responses to increasingly variable environments. The American Naturalist, E57–E64.

Jenouvrier, S., Desprez, M., Fay, R., Barbraud, C., Weimerskirch, H., Delord, K., & Caswell, H. (2018). Climate change and functional traits affect population dynamics of a long-lived seabird. The Journal of Animal Ecology, 87, 906–920.

Jenouvrier, S., Holland, M., Stroeve, J., Barbraud, C., Weimerskirch, H., Serreze, M., & Caswell, H. (2012). Effects of climate change on an emperor penguin population: analysis of coupled demographic and climate models. Global Change Biology, 18, 2756–2770.

Jongejans, E., de Kroon, H., Tuljapurkar, S., & Shea, K. (2010). Plant populations track rather than buffer climate fluctuations. Ecology Letters, 13, 736–743.

Knops, J. M. H., Koenig, W. D., & Carmen, W. J. (2007). Negative correlation does not imply a tradeoff between growth and reproduction in California oaks. Proceedings of the National Academy of Sciences of the United States of America, 104, 16982–16985.

Lawson, C. R., Vindenes, Y., Bailey, L., & van de Pol, M. (2015). Environmental variation and population responses to global change. Ecology Letters, 18, 724–736.

Lenihan, C., & Vuren, D. V. (1996). Growth and survival of juvenile yellow-bellied marmots (*Marmota flaviventris*). Canadian Journal of Zoology, 74, 297–302.

Maldonado-Chaparro, A. A., Blumstein, D. T., Armitage, K. B., & Childs, D. Z. (2018). Transient LTRE analysis reveals the demographic and trait-mediated processes that buffer population growth. Ecology Letters, 23, 1353.

Maldonado-Chaparro, A. A., Read, D. W., & Blumstein, D. T. (2017). Can individual variation in phenotypic plasticity enhance population viability? Ecological Modelling, 352, 19–30.

Marra, P. P., Cohen, E. B., Loss, S. R., Rutter, J. E., & Tonra, C. M. (2015). A call for full annual cycle research in animal ecology. Biology Letters, 11, 20150552.

Merow, C., Dahlgren, J. P., Metcalf, C. J. E., Childs, D. Z., Evans, M. E. K., Jongejans, E., Record, S., Rees, M., Salguero-Gómez, R., & McMahon, S. M. (2014). Advancing population ecology with integral projection models: a practical guide. Methods in Ecology and Evolution, 5, 99–110.

Monclús, R., Pang, B., & Blumstein, D. T. (2014). Yellow-bellied marmots do not compensate for a late start: the role of maternal allocation in shaping life-history trajectories. Evolutionary Ecology, 28, 721–733.

Morgan, I. J., & Metcalfe, N. B. (2001). Deferred costs of compensatory growth after autumnal food shortage in juvenile salmon. Proceedings. Biological Sciences, 268, 295–301.

Morris, W. F., Altmann, J., Brockman, D. K., Cords, M., Fedigan, L. M., Pusey, A. E., Stoinski, T. S., Bronikowski, A. M., Alberts, S. C., & Strier, K. B. (2011). Low demographic variability in wild primate populations: fitness impacts of variation, covariation, and serial correlation in vital rates. The American Naturalist, 177, E14–E28.

Morris, W. F., Pfister, C. A., Tuljapurkar, S., Haridas, C. V., Boggs, C. L., Boyce, M. S., Bruna, E. M., Church, D. R., Coulson, T., Doak, D. F., Forsyth, S., Gaillard, J.-M., Horvitz, C. C., Kalisz, S., Kendall, B. E., Knight, T. M., Lee, C. T., & Menges, E. S. (2008). Longevity can buffer plant and animal populations against changing climatic variability. Ecology, 89, 19–25.

Nouvellet, P., Newman, C., Buesching, C. D., & Macdonald, D. W. (2013). A multi-metric approach to investigate the effects of weather conditions on the demographic of a terrestrial mammal, the european badger (*Meles meles*). PloS One, 8, e68116.

Oli, M. K., & Armitage, K. B. (2004). Yellow-bellied marmot population dynamics: demographic mechanisms of growth and decline. Ecology, 85, 2446–2455.

Orizaola, G., Dahl, E., & Laurila, A. (2014). Compensatory growth strategies are affected by the strength of environmental time constraints in anuran larvae. Oecologia, 174, 131–137.

Ozgul, A., Childs, D. Z., Oli, M. K., Armitage, K. B., & Blumstein, D. T. (2010). Coupled dynamics of body mass and population growth in response to environmental change. Nature, 466, 482–485.

Ozgul, A., Oli, M. K., & Armitage, K. B. (2009). Influence of local demography on asymptotic and transient dynamics of a yellow-bellied marmot metapopulation. The American Naturalist, 173, 517–530.

Ozgul, A., Oli, M. K., Olson, L. E., Blumstein, D. T., & Armitage, K. B. (2007). Spatiotemporal variation in reproductive parameters of yellow-bellied marmots. Oecologia, 154, 95–106.

Paniw, M., Maag, N., Cozzi, G., Clutton-Brock, T., & Ozgul, A. (2019). Life history responses of meerkats to seasonal changes in extreme environments. Science, 363, 631–635.

Paniw, M., Ozgul, A., & Salguero-Gómez, R. (2018). Interactive life-history traits predict sensitivity of plants and animals to temporal autocorrelation. Ecology Letters, 21, 275–286.

Paniw, M., Quintana-Ascencio, P. F., Ojeda, F., & Salguero-Gómez, R. (2017). Accounting for uncertainty in dormant life stages in stochastic demographic models. Oikos, 126, 900–909.

Pfister, C. A. (1998). Patterns of variance in stage-structured populations: evolutionary predictions and ecological implications. Proceedings of the National Academy of Sciences of the United States of America, 95, 213–218.

Picó, F. X., de Kroon, H., & Retana, J. (2002). An extended flowering and fruiting season has few demographic effects in a Mediterranean perennial herb. Ecology, 83, 1991–2004.

Reed, T. E., Grøtan, V., Jenouvrier, S., Sæther, B.-E., & Visser, M. E. (2013). Population growth in a wild bird is buffered against phenological mismatch. Science, 340, 488–491.

Rees, M., & Ellner, S. P. (2009). Integral projection models for populations in temporally varying environments. Ecological Monographs, 79, 575–594.

Robert, A., Bolton, M., Jiguet, F., & Bried, J. (2015). The survival–reproduction association becomes stronger when conditions are good. Proceedings. Biological Sciences, 282, 20151529.

Ruf, T., Bieber, C., Arnold, W., & Millesi, E. (2012). Living in a Seasonal World: Thermoregulatory and Metabolic Adaptations. Springer Science & Business Media.

Rushing, C. S., Hostetler, J. A., Sillett, T. S., Marra, P. P., Rotenberg, J. A., & Ryder, T. B. (2017). Spatial and temporal drivers of avian population dynamics across the annual cycle. Ecology, 98, 2837–2850.

Salguero-Gómez, R., Jones, O. R., & Archer, C. R., et al. (2015). The COMPADRE Plant Matrix Database: an open online repository for plant demography. Journal of Ecology, 103, 202–218.

Salguero-Gómez, R., Jones, O. R., Archer, C. R., et al. (2016). COMADRE: a global data base of animal demography. The Journal of Animal Ecology, 85, 371–384.

Schwartz, O. A., & Armitage, K. B. (2002). Correlations between weather factors and life-history traits of yellow-bellied marmots. Proceedings of the 3rd International Marmot Conference, Cheboksary, Russia.

Schwartz, O. A., & Armitage, K. B. (2005). Weather influences on demography of the yellow-bellied marmot (*Marmota flaviventris*). Journal of Zoology, 265, 73–79.

Schwartz, O. A., Armitage, K. B., & Van Vuren, D. (1998). A 32-year demography of yellow-bellied marmots (*Marmota flaviventris*). Journal of Zoology, 246, 337–346.

Teller, B. J., Adler, P. B., Edwards, C. B., Hooker, G., & Ellner, S. P. (2016). Linking demography with drivers: climate and competition. Methods in Ecology and Evolution, 7, 171–183.

Töpper, J. P., Meineri, E., Olsen, S. L., Rydgren, K., Skarpaas, O., & Vandvik, V. (2018). The devil is in the detail: non-additive and context-dependent plant population responses to increasing temperature and precipitation. Global Change Biology, 24, 4657–4666.

Van de Pol, M., Bailey, L. D., McLean, N., Rijsdijk, L., Lawson, C. R., & Brouwer, L. (2016). Identifying the best climatic predictors in ecology and evolution. Methods in Ecology and Evolution, 7, 1246–1257.

Van de Pol, M., Vindenes, Y., Sæther, B.-E., Engen, S., Ens, B. J., Oosterbeek, K., & Tinbergen, J. M. (2010). Effects of climate change and variability on population dynamics in a long-lived shorebird. Ecology, 91, 1192–1204.

Van Vuren, D. (2001). Predation on yellow-bellied marmots (*Marmota flaviventris*). The American Midland Naturalist, 145, 94–100.

Van Vuren, D., & Armitage, K. B. (1994). Survival of dispersing and philopatric yellow-bellied marmots: what is the cost of dispersal? Oikos, 69, 179–181.

Varpe, Ø. (2017). Life history adaptations to seasonality. Integrative and Comparative Biology, 57, 943–960.

Villellas, J., Doak, D. F., García, M. B., & Morris, W. F. (2015). Demographic compensation among populations: what is it, how does it arise and what are its implications? Ecology Letters, 18, 1139–1152.

Van Vuren, D., & Armitage, K. B. (1991). Duration of snow cover and its influence on life-history variation in yellow-bellied marmots. Canadian Journal of Zoology, 69, 1755–1758.

Williams, J. L., Miller, T. E. X., & Ellner, S. P. (2012). Avoiding unintentional eviction from integral projection models. Ecology, 93, 2008–2014.

Woodroffe, R., Groom, R., & McNutt, J. W. (2017). Hot dogs: High ambient temperatures impact reproductive success in a tropical carnivore. The Journal of Animal Ecology, 86, 1329–1338.

