## Supplementary Material for "Assessing seasonal demographic covariation to understand environmental-change impacts on a hibernating mammal"

### Supporting Material S1 - Bayesian population model

#### *Model description*

The Bayesian model was parameterized for a total of 1,412 female yellow-bellied marmots censuses in eight colonies: bench, boulder, cliff, gothic, marmot meadow, picnic, river, and stonefield. We parameterized the latent variable, $Q_y$, as suggested in Hindle et al. (2018). The standard deviation of $Q_y$ was constrained to 1 and the associated slope parameter, $\beta_q$, for summer survival ($\theta_S$) was constrained to vary positively (SD = 0.2) to make the model identifiable. To ensure identifiability, we also used the sum-to-zero constraint (Kaufman, Sain, & Others, 2010) on the categorical covariate (*i.e.*, stage) in the submodels. This constrains the difference between the model mean of each submodel, $\alpha_0$, and the parameters at each level of a categorical variable, e.g., $\alpha_{a\sigma w}$ [stage], to sum to zero (see Table 1 in main text). In addition, we *z*-transformed the continuous state variable mass ($\mu = 0$; $\sigma = 1$) in all submodels to facilitate convergence of the three chains (Kruschke, 2010).

All model parameters were estimated in JAGS, using the *R* package *jagsUI* (Keller 2015). We used hyperpriors for the random year effects, $\varepsilon_Y$. The hyperpriors were defined as a normal distribution $N(0,\tau)$. We used uninformative, normally distributed priors for all $\alpha$ and $\beta$

parameters (Table 1) with $\mu = 0$ and $1/\sigma^2 = 1\times10^{-6}$, except for the $\beta_q$, where $1/\sigma^2 = 1\times10^{-4}$. Priors for parameters describing variation, *i.e.*, the $\tau$ parameters for the normal likelihood functions and for the hyperpriors describing the random year effect, were defined via a uniform distribution, $1/\tau \sim unif(0,100)$. We ran three chains, each with a burn-in of 1,000,000 iterations. The chains were thinned, selecting every 200$^{th}$ posterior parameter sample after the initial burn-in for a total of 1,000 samples per chain. Convergence after the burn-in was assessed visually and with posterior predictive checks as in Paniw, Quintana-Ascencio, Ojeda, & Salguero-Gómez (2017).

For submodels in which $\beta_q$ was significant (Table S1.1), we also tested for stage-specific differences in demographic responses to environmental quality, *i.e.*, fitting an interaction term, $\beta_{qa}$[stage], between $Q_y$ and stage $a$. None of the interaction terms were significant (95 % credible interval crossed 0), and we therefore kept only the main effects of $Q_y$, described by the $\beta_q$, for further analyses.

*Modelling results*

The posterior parameter distributions from the Bayesian model describing seasonal demographic rates and trait transitions in yellow-bellied marmots are summarized in Table S1.1. Posterior checks suggested the latent variable $Q_y$ accounted for the covariation among the demographic processes. That is, including $Q_y$ in the Bayesian model effectively eliminated correlations among the submodel-specific estimates of the random-year effects, $\varepsilon_Y$ (Fig. S1.1). This suggests that for our study, a single latent parameter is sufficient to account for the covariation among the demographic processes (Hindle et al., 2018). Derived values of $Q_y$ for each of the 40 study years were not autocorrelated and showed no trend (Fig. S1.2).

**Table S1.1** Summary of posterior distributions for parameters in Bayesian model describing the demographic processes of yellow-bellied marmots (see Table 1 in the main text for parameter names). Posterior means in bold identify parameters with 95% credible intervals not crossing 0. $\bar{R}$- Gelman-Rubin convergence diagnostic (at convergence, $\bar{R}$= 1); n.eff – effective sample size; f - proportion of the posterior with the same sign as the mean.

| | | mean | sd | 2.5 % | 50 % | 97.5 % | $\bar{R}$ | n.eff | f |
|---|---|---|---|---|---|---|---|---|---|
| Winter survival | $\alpha_{0\theta W}$ | **0.72** | 0.11 | 0.51 | 0.72 | 0.94 | 1.00 | 2673 | 1.00 |
| | $\beta_{z\theta W}$ | **0.59** | 0.14 | 0.32 | 0.59 | 0.87 | 1.00 | 3000 | 1.00 |
| | $\beta_{q\theta W}$ | **0.27** | 0.13 | 0.04 | 0.26 | 0.55 | 1.00 | 3000 | 1.00 |
| | $\alpha_{a\theta W}$[J] | -0.28 | 0.20 | -0.69 | -0.28 | 0.10 | 1.00 | 3000 | 0.93 |
| | $\alpha_{a\theta W}$[Y] | **0.31** | 0.11 | 0.09 | 0.30 | 0.52 | 1.00 | 3000 | 1.00 |
| | $\alpha_{a\theta W}$[N] | 0.00 | 0.15 | -0.28 | 0.00 | 0.30 | 1.00 | 1699 | 0.51 |
| | $\alpha_{a\theta W}$[R] | -0.02 | 0.12 | -0.25 | -0.02 | 0.22 | 1.00 | 3000 | 0.56 |
| | $\tau_Y$ | 0.37 | 0.10 | 0.13 | 0.38 | 0.57 | 1.00 | 3000 | 1.00 |
| Mass next (W) | $\alpha_{0z*W}$ | **4.79** | 0.32 | 4.14 | 4.78 | 5.43 | 1.00 | 1786 | 1.00 |
| | $\beta_{zz*W}$ | **0.59** | 0.02 | 0.55 | 0.59 | 0.63 | 1.00 | 1957 | 1.00 |
| | $\beta_{qz*W}$ | **0.28** | 0.12 | 0.06 | 0.28 | 0.49 | 1.00 | 3000 | 0.99 |
| | $\beta_{zaz*W}$[J] | **0.13** | 0.03 | 0.08 | 0.13 | 0.19 | 1.00 | 2319 | 1.00 |
| | $\beta_{zaz*W}$[Y] | **-0.09** | 0.04 | -0.17 | -0.09 | -0.01 | 1.00 | 3000 | 0.99 |
| | $\beta_{zaz*W}$[N] | -0.04 | 0.04 | -0.12 | -0.04 | 0.05 | 1.00 | 2438 | 0.81 |
| | $\beta_{zaz*W}$[R] | 0.00 | 0.03 | -0.06 | 0.00 | 0.06 | 1.00 | 3000 | 0.55 |
| | $\alpha_{az*W}$[J] | **-1.75** | 0.36 | -2.45 | -1.76 | -1.03 | 1.00 | 1951 | 1.00 |
| | $\alpha_{az*W}$[Y] | 1.12 | 0.57 | -0.03 | 1.13 | 2.22 | 1.00 | 3000 | 0.97 |
| | $\alpha_{az*W}$[N] | 0.43 | 0.64 | -0.83 | 0.43 | 1.70 | 1.00 | 2382 | 0.75 |
| | $\alpha_{az*W}$[R] | 0.20 | 0.46 | -0.69 | 0.21 | 1.05 | 1.00 | 3000 | 0.67 |
| | $\tau_{z*W}$ | **0.48** | 0.01 | 0.46 | 0.48 | 0.50 | 1.00 | 1377 | 1.00 |
| | $\tau_Y$ | 0.29 | 0.10 | 0.05 | 0.31 | 0.45 | 1.01 | 2431 | 1.00 |
| Transition to reproductive adult | $\alpha_{0\varpi 0}$ | **0.91** | 0.21 | 0.51 | 0.91 | 1.33 | 1.00 | 1033 | 1.00 |
| | $\beta_{z\varpi 0}$ | **0.46** | 0.12 | 0.23 | 0.46 | 0.70 | 1.00 | 3000 | 1.00 |
| | $\beta_{q\varphi 0}$ | **0.69** | 0.34 | 0.04 | 0.68 | 1.37 | 1.00 | 3000 | 0.98 |
| | $\alpha_{a\varphi 0}$[Y] | **-0.74** | 0.15 | -1.03 | -0.74 | -0.43 | 1.00 | 3000 | 1.00 |
| | $\alpha_{a\varpi 0}$[N] | -0.12 | 0.16 | -0.42 | -0.12 | 0.19 | 1.00 | 3000 | 0.79 |
| | $\alpha_{a\varphi 0}$[R] | **0.86** | 0.13 | 0.60 | 0.85 | 1.13 | 1.00 | 3000 | 1.00 |
| | $\tau_Y$ | 0.83 | 0.29 | 0.13 | 0.86 | 1.35 | 1.00 | 2620 | 1.00 |
| Summer survival | $\alpha_{0\theta S}$ | **2.10** | 0.14 | 1.83 | 2.09 | 2.39 | 1.00 | 3000 | 1.00 |
| | $\beta_{z\theta S}$ | 0.01 | 0.17 | -0.32 | 0.01 | 0.36 | 1.00 | 3000 | 0.52 |
| | $\beta_{q\theta S}$ | 0.07 | 0.24 | -0.40 | 0.07 | 0.52 | 1.00 | 1529 | 0.61 |
| | $\alpha_{a\theta S}$[Y] | **-0.94** | 0.24 | -1.39 | -0.94 | -0.46 | 1.00 | 3000 | 1.00 |
| | $\alpha_{a\theta S}$[N] | **-0.40** | 0.17 | -0.74 | -0.39 | -0.05 | 1.00 | 3000 | 0.99 |
| | $\alpha_{a\theta S}$[R] | **1.33** | 0.19 | 0.96 | 1.33 | 1.71 | 1.00 | 1693 | 1.00 |
| | $\tau_Y$ | 0.54 | 0.14 | 0.27 | 0.53 | 0.84 | 1.00 | 2007 | 1.00 |
| Mass next (S) | $\alpha_{0z*S}$ | **6.46** | 0.24 | 5.99 | 6.46 | 6.93 | 1.00 | 2404 | 1.00 |
| | $\beta_{zz*S}$ | **0.65** | 0.02 | 0.61 | 0.65 | 0.69 | 1.00 | 2281 | 1.00 |
| | $\beta_{qz*S}$ | -0.07 | 0.07 | -0.20 | -0.07 | 0.08 | 1.00 | 2211 | 0.83 |
| | $\beta_{zaz*S}$[Y] | **-0.15** | 0.02 | -0.19 | -0.15 | -0.11 | 1.00 | 3000 | 1.00 |
| | $\beta_{zaz*S}$[N] | -0.02 | 0.03 | -0.07 | -0.02 | 0.04 | 1.00 | 3000 | 0.76 |
| | $\beta_{zaz*S}$[R] | **0.17** | 0.02 | 0.12 | 0.17 | 0.21 | 1.00 | 3000 | 1.00 |
| | $\alpha_{az*S}$[Y] | **2.26** | 0.26 | 1.77 | 2.25 | 2.75 | 1.00 | 3000 | 1.00 |
| | $\alpha_{az*S}$[N] | 0.32 | 0.36 | -0.39 | 0.32 | 1.03 | 1.00 | 3000 | 0.82 |
| | $\alpha_{az*S}$[R] | **-2.58** | 0.30 | -3.15 | -2.57 | -1.99 | 1.00 | 3000 | 1.00 |
| | $\tau_{z*S}$ | **0.42** | 0.01 | 0.40 | 0.42 | 0.44 | 1.00 | 3000 | 1.00 |
| | $\tau_Y$ | 0.27 | 0.04 | 0.20 | 0.27 | 0.35 | 1.00 | 3000 | 1.00 |
| # recruits | $\alpha_{0\varphi 1}$ | **1.45** | 0.04 | 1.37 | 1.45 | 1.52 | 1.00 | 3000 | 1.00 |
| | $\beta_{z\varphi 1}$ | **0.11** | 0.02 | 0.06 | 0.11 | 0.16 | 1.00 | 2387 | 1.00 |
| | $\beta_{q\varphi 1}$ | 0.00 | 0.07 | -0.13 | 0.00 | 0.14 | 1.00 | 1460 | 0.53 |
| | $\tau_Y$ | 0.15 | 0.04 | 0.09 | 0.16 | 0.24 | 1.00 | 3000 | 1.00 |
| Juvenile mass | $\alpha_{0z_j^*}$ | **4.53** | 0.54 | 3.48 | 4.52 | 5.55 | 1.00 | 3000 | 1.00 |
| | $\beta_{zz_j^*}$ | **0.45** | 0.04 | 0.37 | 0.45 | 0.52 | 1.00 | 3000 | 1.00 |
| | $\beta_{qz_j^*}$ | 0.17 | 0.11 | -0.05 | 0.17 | 0.39 | 1.00 | 2915 | 0.93 |
| | $\tau_{z_j^*}$ | **0.73** | 0.02 | 0.70 | 0.73 | 0.76 | 1.00 | 3000 | 1.00 |
| | $\tau_Y$ | 0.32 | 0.07 | 0.18 | 0.32 | 0.45 | 1.00 | 1042 | 1.00 |

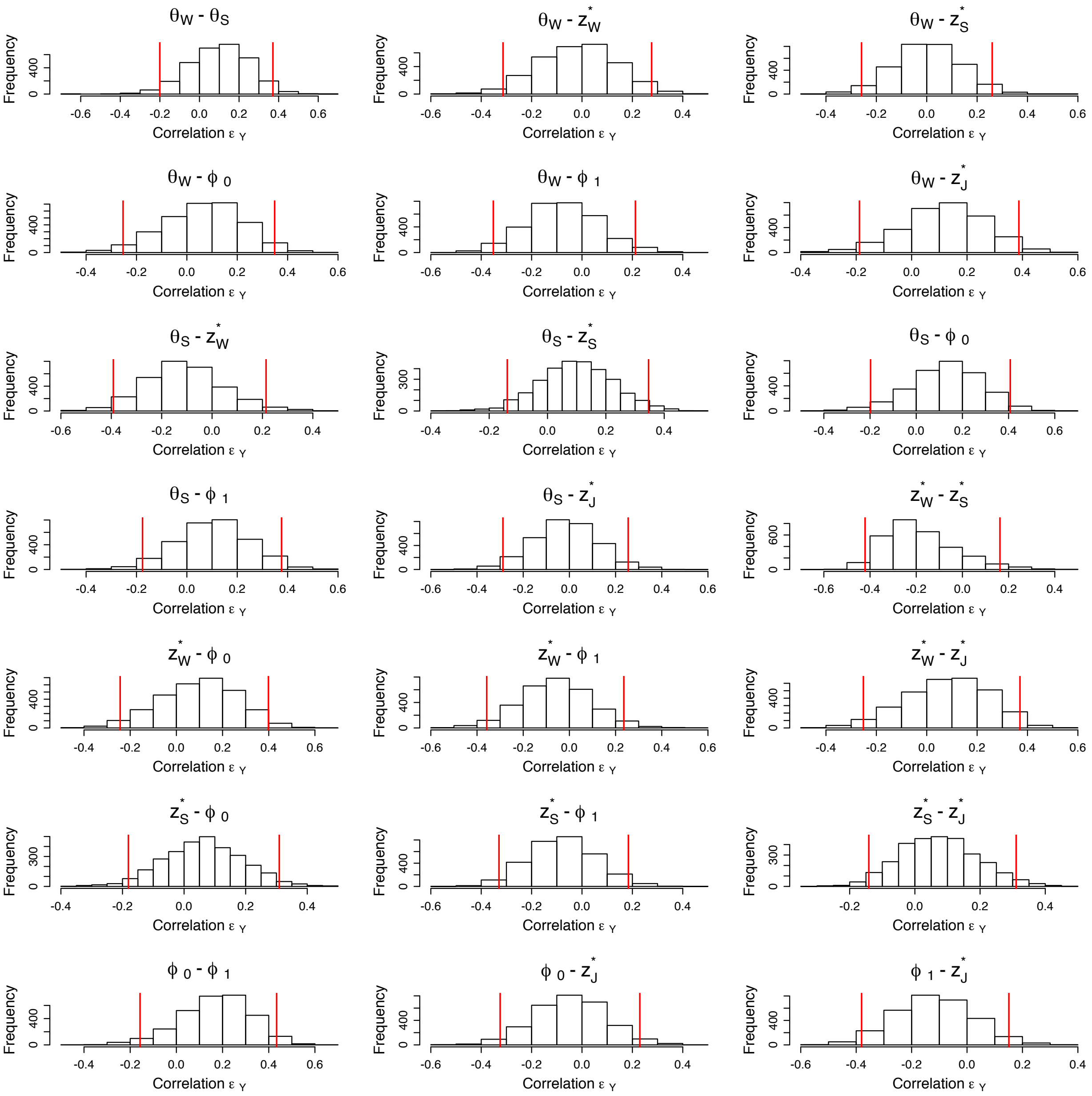

**Figure S1.1:** Correlation between the submodel-specific year effects ($\varepsilon_Y$) in the yellow-bellied marmot Bayesian model. Correlations were calculated for each of 3,000 posterior parameters values of each $\varepsilon_Y$. Vertical red lines show the 95 % credible interval of the distribution of the correlations. The submodels include winter ($\theta_W$) and summer ($\theta_S$) survival, winter ($z_W^*$) and summer ($z_S^*$) mass change, probability of reproducing ($\varphi_0$), number of recruits ($\varphi_1$), and juvenile mass ($z_J^*$).

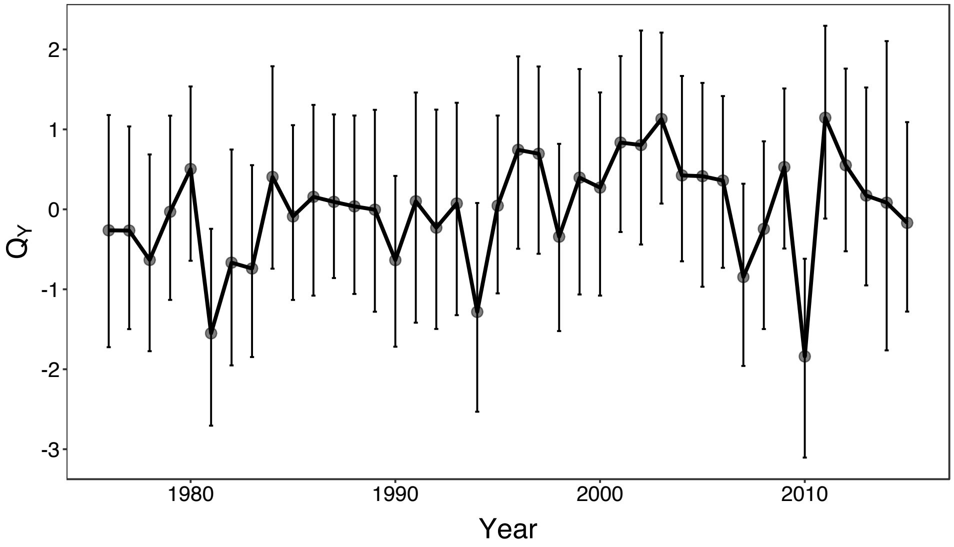


**Figure S1.2:** Distribution of year-specific values of $Q_y$ derived from the Bayesian model. Points and error bars show means and 95 % credible intervals of values obtained from 3,000 samples of the posterior parameter distribution. Visual inspection (using the *acf* function in R) indicated a lack of autocorrelation in the distribution of $Q_y$. A Box-Pierce test ($X^2$= 0.07, d.f. = 1, $p$ = 0.79) and Jarque-Bera test ($X^2$= 4.5, d.f. = 2, $p$ = 0.10) further suggest that $Q_y$ are independent and normally distributed, respectively.

The latent variable $Q_y$ was correlated with several environmental variable measured at the study site 1976-2013 (Fig. S1.3). To explore these relationships further, we first accounted for the collinearity among environmental variables by performing a principle-component analysis (PCA) on all environmental variables that were correlated with $Q_y$ (*i.e.*, 95 % CI did not overlap 0 in Fig. S1.3). We used the R package *psych* (Revell, 2008) to perform the PCA on rescaled data to mean = 0 and SD = 1. The PCA reduced the dimensionality of the environmental data by quantifying two main axes (associated eigenvalues > 1; Legendre & Legendre 2012) of environmental conditions that together explained 86% of the variation in the environmental data (Fig. S1.4a). PCA axis 1 largely captured among-year variation in summer length vs. winter length and snow cover, while PCA 2 captured variation in winter and spring temperature severity. We then modeled average derived $Q_y$ (1976-2013) as a function of PCA 1 and PCA 2, and their interaction, using a generalized additive model (GAM) to flexibly model non-linear interactions. We fitted the GAM using the *mgcv* package in R (Wood 2006). In parameterizing the GAM, we used tensor product smooth terms (*te*), with thin-plate regression splines as the marginal bases for the effects of the PCA axes. When applying smooths, we used four-knot locations. We also applied a degrees-of-freedom inflation factor (*gamma*) of 1.4 to avoid overfitting due to overly flexible smooths. In addition to PCA 1 and PCA 2, we tested for the effect of population density (and density of adults) at the beginning of a year on $Q_y$ by modeling interactions between density and the PCA axes, but AIC comparisons indicated that including density did not improve model fit (differences in AIC < 2). The most parsimonious GAM took the following form:

$$Q_y \sim te(\text{PCA 1, PCA 2}_{\text{df:3}}),$$

where *df* represents the amount of nonlinearity in the model component, with *df*=1 indicating linear fit. Results from the GAM indicate that the two axes of environmental conditions explain 45.7 % of the variance in $Q_y$, which further supports the conlcusion that $Q_y$ is an aggregate measure of the partially unobserved quality of the environment experienced by yellow-bellied marmots (Fig. S1.4b) (Hindle et al., 2018).

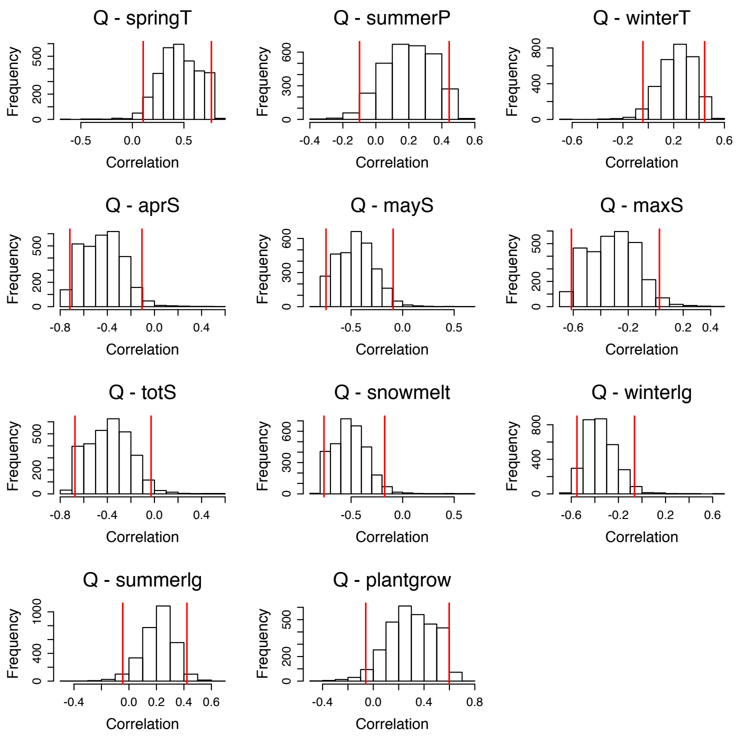


**Figure S1.3:** Correlation between the latent variable $Q_y$, quantifying the overall environmental quality in the yellow-bellied marmot Bayesian model, and environmental variables measured at the study site (across all eight colonies) since 1976. Correlations were calculated for each of 3,000 posterior parameters values of $Q_y$. From top left to bottom right, the environmental variables depict: springT, mean of the average daily temperature in spring (April-May); summerP, mean of daily precipitation in summer (June-September); winterT, mean of the average daily temperature in winter (October-March); aprS, snow depth (inches) on t April 1st; mayS, snow depth (inches) on May 1st; maxS, maximum snow depth (inches) during the winter; totS, total snowfall during the winter; snowmelt, Julian date of snowmelt (when no snow remained on the ground at the weather station); winterlg, winter length (number of days from the first snowfall the previous year to the last snowfall of the current year); summerlg, summer length (number of days between the last snowfall in spring and the first snowfall in the fall ); and plantgrow, length of the growing season (number of days between the day of 1st bare ground to the 1st day with average  mean temperature below 0°C). Details on the environmental variables can be found in (O. A. Schwartz & Armitage, 2002; Orlando A. Schwartz & Armitage, 2005).

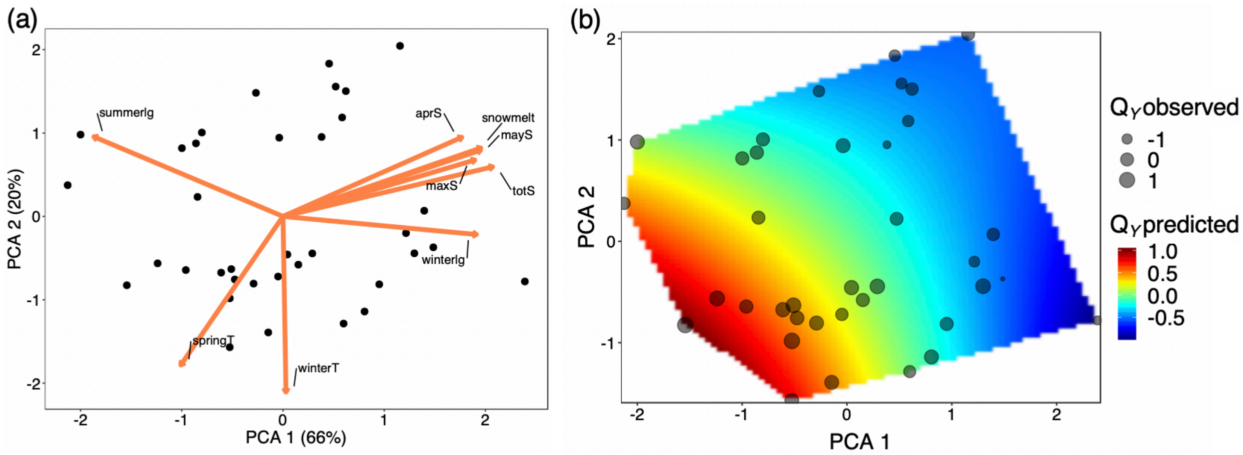


**Figure S1.4:** Two main PCA axes explain the majority of the variation in environmental variables measured at the study site (a) and predict the latent variable $Q_y$, quantifying the overall environmental quality in the yellow-bellied marmot Bayesian model. In (a), points depict 37 years (1976-2012). Arrow lengths are proportional to the loadings of each environmental variable onto the two axes. The environmental variables depict: springT, mean of the average daily temperature in spring (April-May); winterT, mean of the average daily temperature in winter (October-March); aprS, snow depth (inches) on t April 1st; mayS, snow depth (inches) on May 1st; maxS, maximum snow depth (inches) during the winter; totS, total snowfall during the winter; snowmelt, Julian date of snowmelt (when no snow remained on the ground at the weather station); winterlg, winter length (number of days from the first snowfall the previous year to the last snowfall of the current year); and summerlg, summer length (number of days between the last snowfall in spring and the first snowfall in the fall). In (b), the raster plot shows predicted $Q_y$ values across the two PCA axes, while points indicate 'observed' $Q_y$ (*i.e.*, derived from the Bayesian model). Predictions are limited to the range of observed PCA scores.

Predictions from the most parsimonious demographic-process models (*i.e.*, demographic rates and trait transitions) showed that all processes increased with body mass; but body mass played the least significant role in summer survival ($\theta_S$; Fig. S1.5). Reproductive adults also had the highest mean survival, trait transitions, and reproduction; they lost and gained relatively little mass the heavier they were in the winter and summer season, respectively (Fig. S1.5). The environmental quality, $Q_y$, had a positive, additive effect on most demographic processes (although the 95 % C.I. of this effect did not cross 0 only for winter demographic processes), except for summer mass gain ($z_S^*$) and recruitment ($\varphi_1$), where the effect was slightly negative but not significant (Table S1.1).

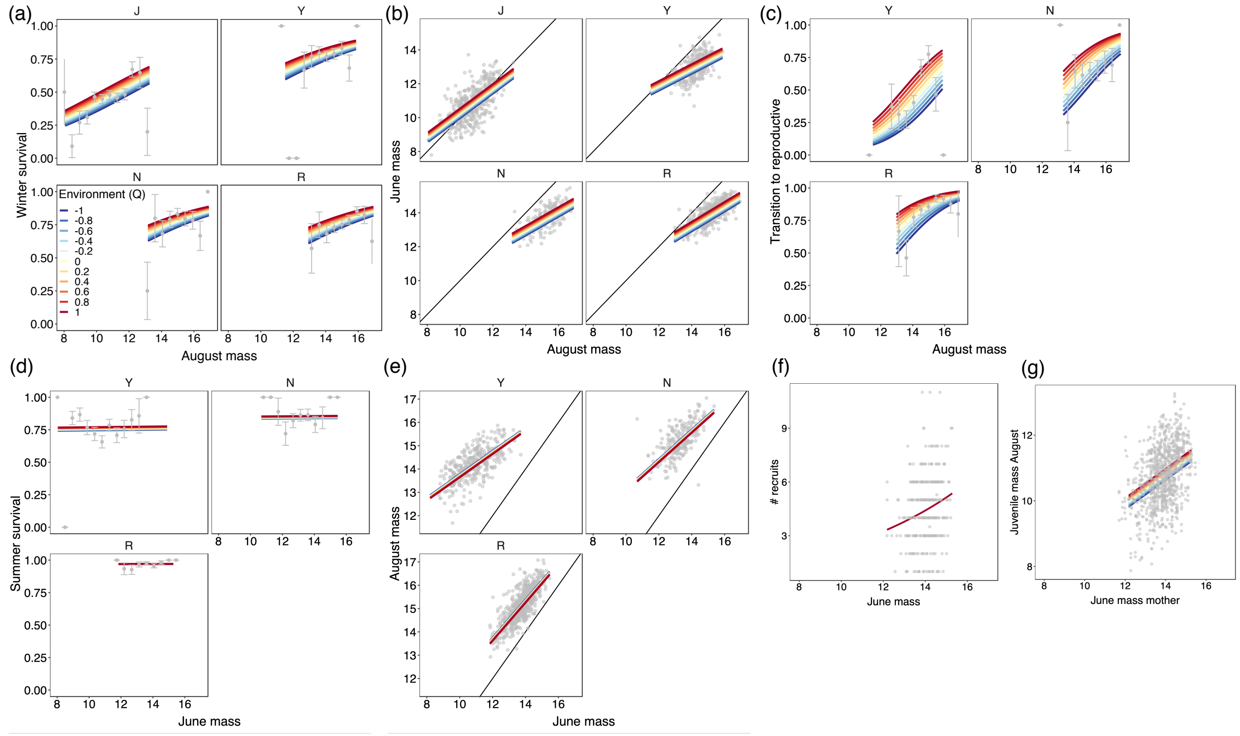


**Figure S1.5** Predictions (lines) from the most parsimonious submodels within the Bayesian framework describing seasonal demographic rates and trait transitions of yellow-bellied marmots. Predictions are plotted for the minimum and maximum observed cube-root transformed masses per stage. Stages include juveniles (J), yearlings (Y), non-reproductive adults (N), and reproductive adults (R). Black lines depict stable masses (intercept = 0; slope =1 for mass change). Points show observed values. For binomial responses in (a), (c), and (d) points and error bars show mean ± S.E., respectively, of observed values across different mass classes.

$Q_y$ also showed a nonlinear relationship with annual changes in population size observed over the 40 years of the study period (Fig. S1.6). That is, changes in population size were positively related to $Q_y$ when $Q_y$ approached large values but were little affected at $Q_y$ close to 0. At the same time, neither individual environment variables nor the main environmental axes defined in a PCA (see above) were correlated with changes in population size (Fig. S1.7).

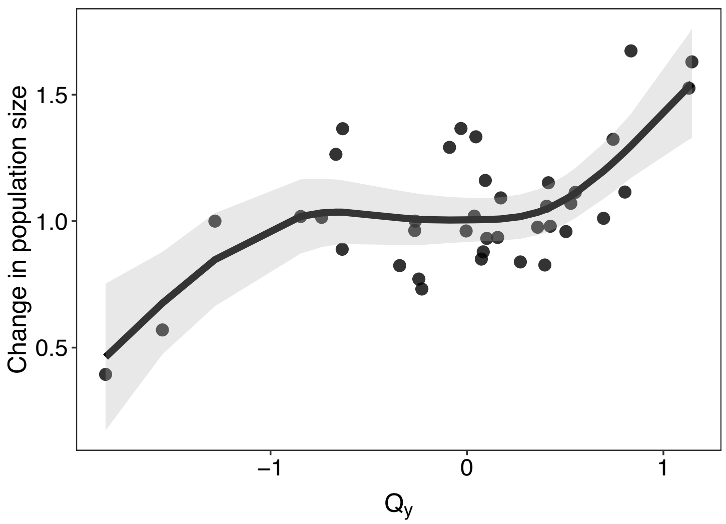


**Figure S1.6:** Observed changes in population size (*i.e.*, annual realized growth rate in the empirical data on yellow-bellied marmots) modeled as a function of average posterior values of $Q_y$, quantifying the overall environmental quality in the Bayesian model. A GAM model very similar in structure to the one described in Fig. S1.4 was fit to the data. Points show observed values while the line and the shaded area show averaged predicted values and the 95 % prediction interval, respectively.

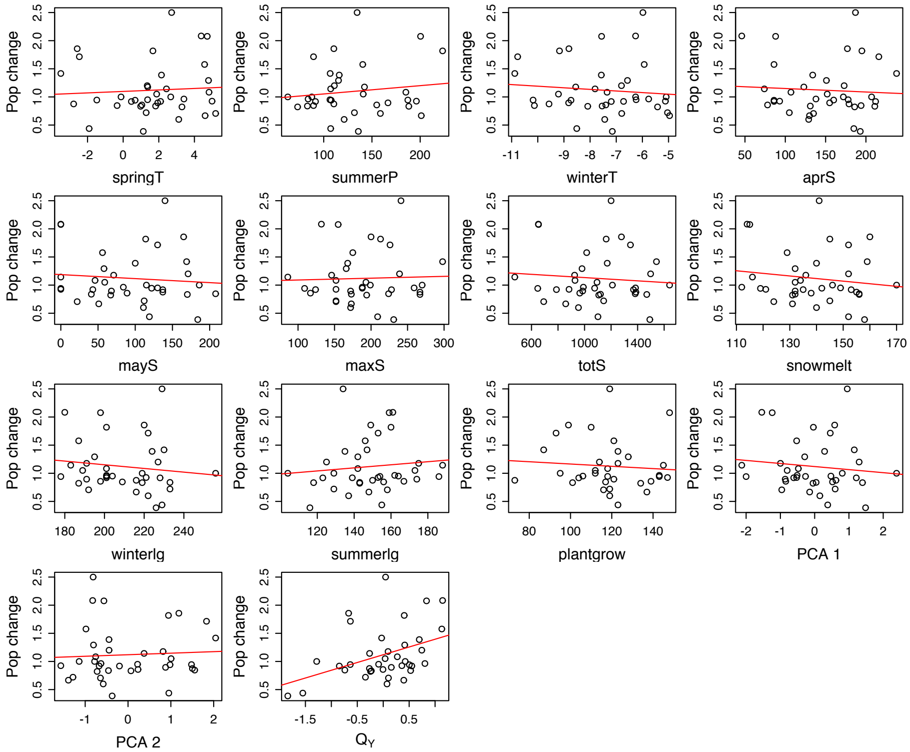


**Figure S1.7:** Observed changes in population size (*i.e.*, annual realized growth rate in the empirical data on yellow-bellied marmots) only correlate with average posterior values of $Q_y$, quantifying the overall environmental quality in the Bayesian model. Points show observed values 1976-2013. Lines depicts best-fit predictions from a simple regression model. The slope of this model was only significant in the case of $Q_y$ (Wald test, slope = 0.27, $p$ = 0.01).

**Supporting Material S2 - Integral projection model**

Using the most parsimonious GLMMs from the Bayesian model, we constructed a stage-mass classified, stochastic Integral Projection Model (IPM) for the winter (W) and summer (S) season. Survival and trait change of each of the four life-history stages (*i.e.*, juveniles, yearlings, non-reproductive adults, and reproductive adults) and recruitment were described by IPM kernels (corresponding to survival-growth [P] and recruitment [F] kernels; see Ellner et al., 2016). Upon survival and trait transitions, individuals then transitioned to, or stayed in, a given stage (see Fig. 1 in the main text). At the beginning of the summer season ($t$+1), number of yearling ($n_Y$) in the mass range $z'$ = [$z'$, $z'$+d$z$] in the populations was therefore characterized by number of juveniles ($n_J$) in the mass range $z$ = [$z$, $z$+d$z$] in winter ($t$), which survived ($\theta$) and changed mass (shrank) ($\gamma$) under a season-specific environmental quality, Q:

$$n_Y(z',t+1) = \int_{\Omega} [\theta_W^J(z,Q,t)\,\gamma_W^J(z,z',Q,t)]n_J(z,t)\mathrm{d}z$$

Following the example above, the number of reproductive adults (R) at the beginning of the summer season was comprised of stages $a$ = juveniles, non-reproductive, or reproductive adults, in which individuals survived, changed mass, and reproduced with a probability = $\varphi_0$:

$$n_R(z',t+1) = \sum_{a\neq J} \int_{\Omega} [\theta_W^a(z,Q,t)\,\gamma_W^a(z,Q,t)\,\varphi_{0W}^a(z,Q,t)]n_a(z,t)\mathrm{d}z$$

The number of non-reproductive adults (N) at the beginning of the summer season was then defined as:

$$n_N(z',t+1) = \sum_{a\neq J} \int_{\Omega} [\theta_W^a(z,Q,t)\,\gamma_W^a(z,z',Q,t)\,(1-\varphi_{0W}^a(z,Q,t))]n_a(z,t)\mathrm{d}z$$

Throughout the summer season, individuals of all stages $a$ with the exception of juveniles survived and gained mass until the beginning of the winter season, described as:

$$n_{a\neq J}(z', t+1) = \int_{\Omega} [\theta_S^{a\neq J}(z, Q, t)\, \gamma_S^{a\neq J}(z, z', Q, t)] n_{a\neq J}(z, t) \mathrm{d}z$$

The number of juveniles (J) at the beginning of the winter season was the result of the reproductive effort of adults (R; see main text). Juveniles had a likelihood of being in a given mass class defined by $\varphi_2$:

$$n_J(z', t+1) = \int_{\Omega} [\theta_S^{R}(z, Q, t)\, \varphi_1(z, Q, t)/2\, \varphi_2(z, Q, t)] n_R(z, t) \mathrm{d}z$$

We obtained annual population dynamics from the periodic product of the winter and summer IPMs. Although we corrected for eviction in the IPMs, potential eviction was minimal and largest for recruits (Fig. S2.1).

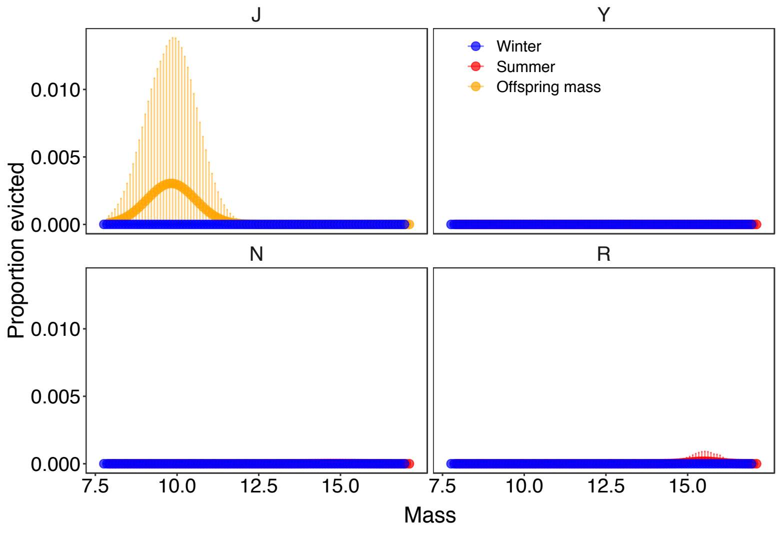


**Figure S2.1:** Proportion of individuals in each stage that would have been evicted without correcting for eviction in the IPM. This proportion was calculated by comparing the proportion of individuals surviving and changing mass between a model that accounted for eviction in survival-growth kernels and one that did not.

We assessed the robustness and accuracy of the annual IPM by calculating the correlation between observed stage-specific relative changes in abundance (*i.e.*, annual realized growth rate) and ones obtained from IPM simulations. To obtain abundances from IPM simulations, we iterated seasonal population dynamics from 1976 to 2016, beginning with the population vector $n_a(z)$ of female marmots in different stages $a$ and mass $z$ in the beginning of the winter season 1976, and obtained annual relative changes in $n_a(z)$, *i.e.*, $n_{at+1}(z)/n_{at}(z)$. We used year-specific values of $Q_y$ and $\varepsilon_Y$ obtained from the Bayesian model to construct IPMs at each iteration. To account for parameter uncertainty, we repeated the 40 years of iterations of population dynamics for 1,000 Bayesian model parameters randomly sampled from the posterior distribution.

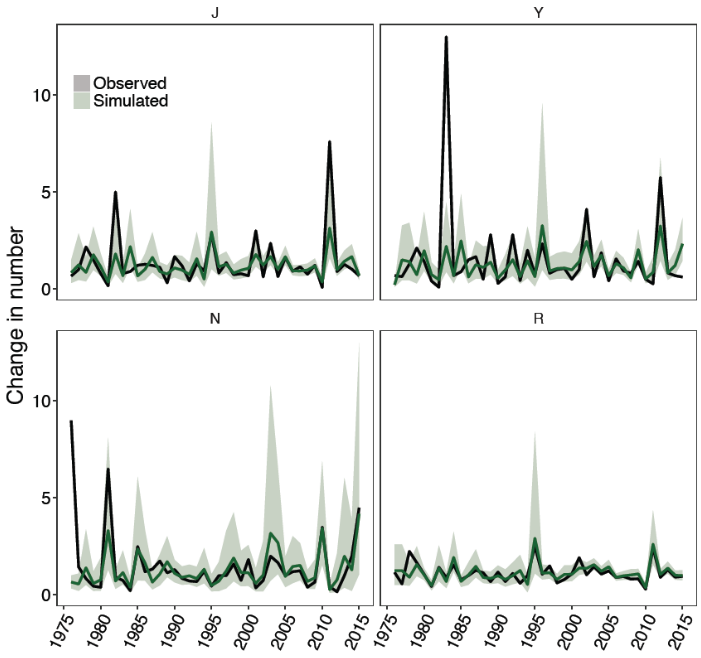


**Figure S2.2:** Relative changes in stage-specific abundances (*i.e.*, annual realized growth rate) observed in the empirical data on yellow-bellied marmots and obtained from IPM simulations. Stages include juveniles (J), yearlings (Y), non-reproductive adults (N), and reproductive adults (R). Shaded area shows the 95 % credible interval of simulated results based on simulations for 1,000 posterior parameters from the Bayesian population model.

Our results show a good match between observed and simulated data (Figs. S2.2 & S2.3). Although at one occasion, we failed to simulate a large increase in the number of juveniles, our simulations closely followed relative increases and decreases in adult abundances, for both non-reproductive (N) and reproductive (R) adults (Fig. S.2.2), with demographic processes of adults contributing most to population dynamics (Supporting Material S3).

Our population model assumes that body mass provides a cumulative summary of an individual marmot's past experience, both social (reproductive stage) and environmental (via $Q_y$). This summary directly affects vital rates (*i.e.*, survival, growth, and reproduction); and the mass distribution within the population, propagated from one season to the next, mediates population responses to future environmental conditions. We made this assumption because endogenous tradeoffs have thus far not been detected in marmots. As all demographic rates covary positively, our study indicates that environmental drivers, along with social aspects of group living primarily affect demography.

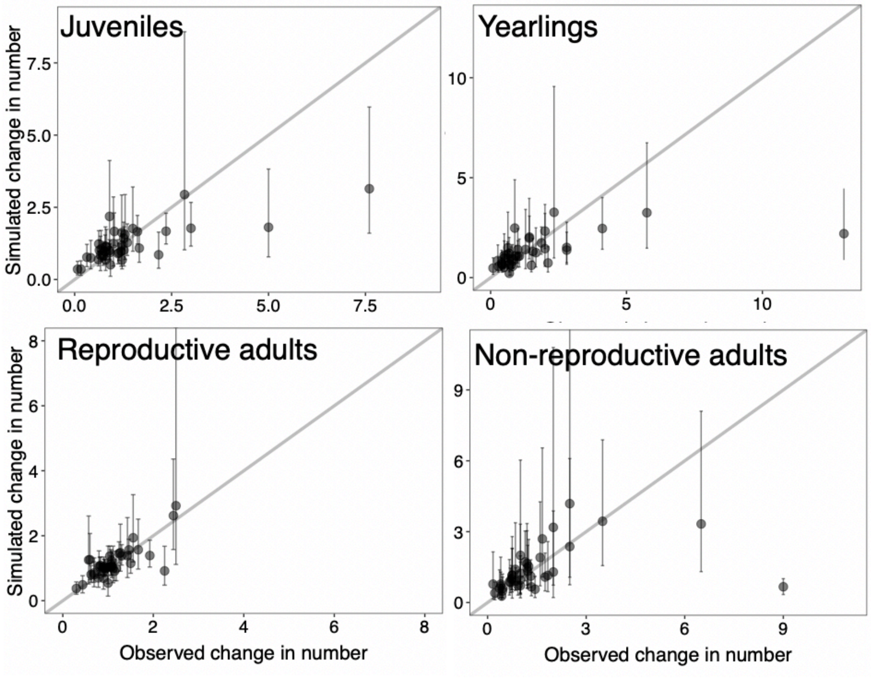


**Figure S2.3:** Relative changes in stage-specific abundances (*i.e.*, annual realized growth rate) obtained from IPM simulations are within the range of observed changes in the empirical data on yellow-bellied marmots except when changes in juveniles, yearlings, and non-reproductive adults are extreme. Error bars show the 95 % credible interval of simulated results and are based on performing simulations for 1,000 posterior parameters from the Bayesian population model. Grey lines depict best-fit lines (intercept = 0; slope =1).

**Supporting Material S3 - Elasticity of the stochastic growth rate to demographic processes**

To calculate the stochastic population growth rate, log $\lambda s$, the simulation of 100,000 years started with the population of individuals with the standing stage and mass distribution at the beginning of the winter season in 2016. At each run of the simulation, a seasonal IPM was constructed with resampled, year-specific $Q_y$ and $\varepsilon_Y$ values. We then calculated the periodic IPM product of the winter and summer IPMs, describing the demographic and trait transitions from the beginning of one winter season to the next (Caswell, 2001; chapter 13). Lastly, we obtained log$\lambda s$ as the mean of the logs of annual realized population growth rates, $\lambda = n_{t+1}/n_t$, where $n_t$ and $n_{t+1}$ are the total number of individuals in the population at time (year) *t* and *t*+1, respectively.

We calculated elasticities of the stochastic growth rate to changes in the mean ($e_S^{\mu}$) and standard deviation ($e_S^{\sigma}$) of mass- and stage-specific demographic rates and trait transitions (*i.e.*, demographic processes) by perturbing a demographic-process function ($\rho$) for a given stage over an interval of masses [*z*, *z*+d*z*] by its mean and standard deviation obtained across 40 years of observed dynamics. Observed dynamics were obtained by resampling year-specific values of $Q_y$ and $\varepsilon_Y$ for 1976-2016 from the Bayesian model. We then built a winter or summer IPM using the perturbed vital rate and obtained a new, perturbed periodic IPM product (*i.e.*, the perturbation kernel). Lastly, we integrated this product into eq. 7.5.2 in Ellner *et al.* (2016) in order to estimate $e_S^{\mu}$ and $e_S^{\sigma}$ from the simulations of population dynamics for T = 90,000 years (*i.e.*, discarding the first 10, 000 years of simulations to ensure no effect of transient population fluctuations):

$$e = E[\frac{\langle v_{t+1}, C_t w_t \rangle}{\langle v_{t+1}, K_t w_t \rangle}],$$

where $C_t$ is the perturbation kernel, $K_t$ is the unperturbed annual IPM kernel at each simulation iteration $t$, $v$ and $w$ are the left and right eigenvectors associated with $K$ at $t$ +1 and $t$, respectively. For instance, if average winter survival in a given mass interval (*i.e.*, IPM bin) for reproductive individuals from 1976-2016 was 0.95, each winter IPM during T simulations was perturbed by subtracting a focal value of winter survival from its 40-year mean; and $e_S^{\mu}$ was calculated by integrating this perturbed IPM as $C_t$ into the above equation. We used an interval of 50 classes, *i.e.*, at 50 of the 100 IPM bins evenly spread from the lowest to highest mass for each stage, when performing the perturbations at the mean posterior parameter values and an interval of 10 classes when performing perturbations across 100 samples posterior parameter samples. Using 50 or 10 mass classes instead of the full 100 allowed us to save computational time for the perturbations, which was increased particularly when we accounted for parameter uncertainty when calculating the elasticities.

We then used the chain rule to assess how this perturbation affected the annual integral projection model (IPM) (see main text) (Haridas & Tuljapurkar, 2005; Rees & Ellner, 2009), and how this perturbed IPM affected log$\lambda_S$- as suggested in (2016). When perturbing trait transitions, we accounted for the non-independence in the transition kernels (*i.e.*, each column of the discretized trait transition kernel had to sum to 1) using *proportional compensation* as suggested by (2017).

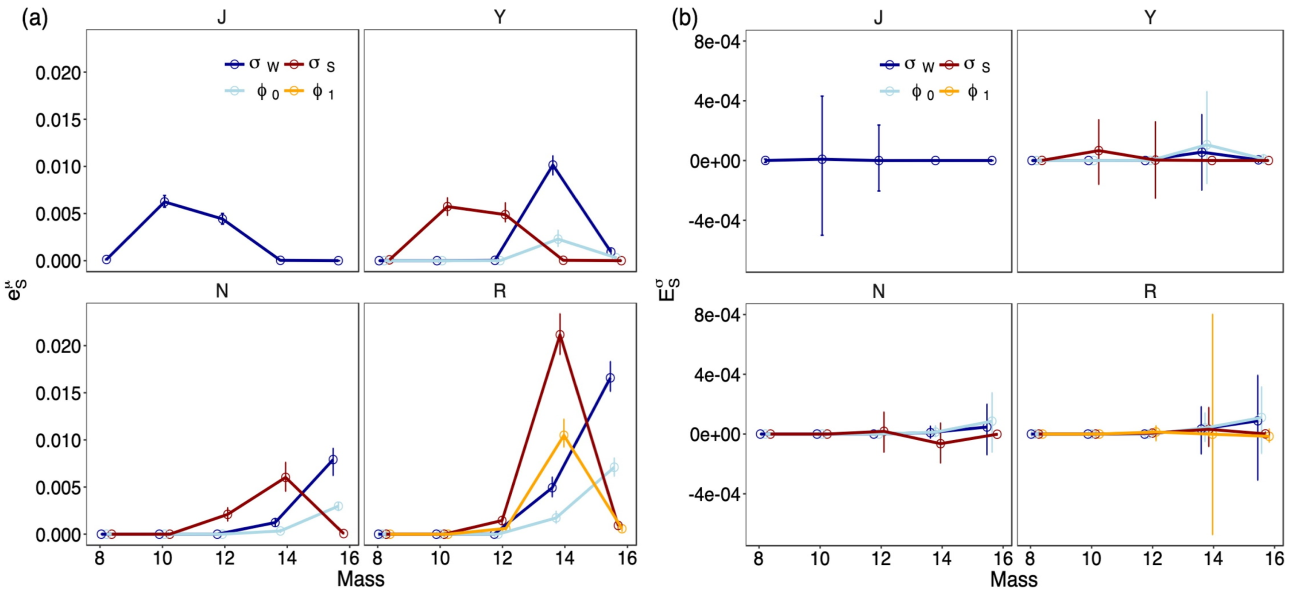


**Figure S3.1:** Elasticities (e) of log λs to changes in (a) the mean ($\mu$) and (b) standard deviation (σ) of stage-specific demographic rates at 10 different mass classes. Demographic rates include winter ($\theta_W$) and summer survival ($\theta_S$), probability of reproducing ($\varphi_0$), and recruitment ($\varphi_1$). Stages are juveniles (J), yearlings (Y), non-reproductive adults (N), and reproductive adults (R). Points and error bars show means and 95 % credible intervals of the elasticities obtained from 100 samples of the posterior parameter distribution.

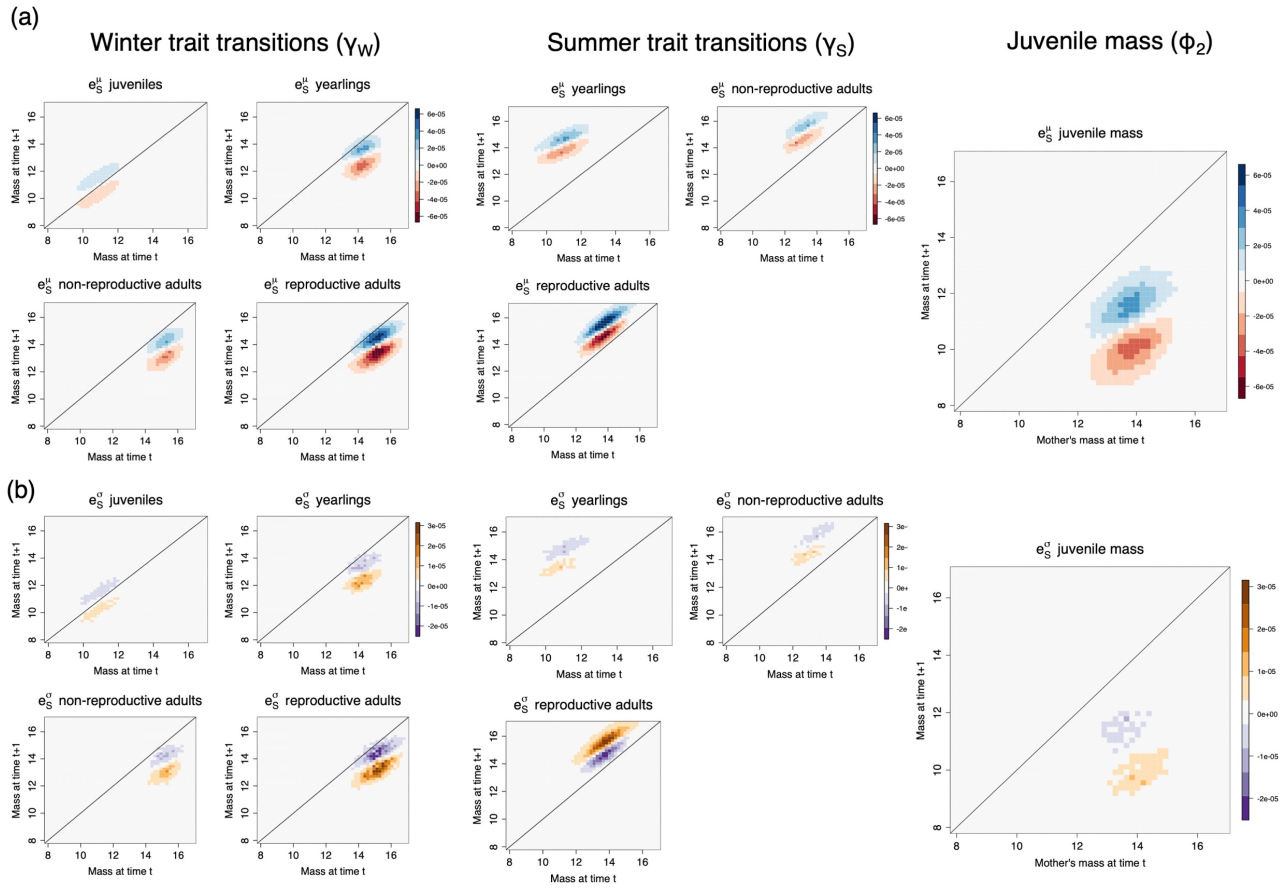


**Figure S3.2:** Elasticities (es) of log λs to changes in (a) the mean (*μ*) and (b) standard deviation (σ) of stage-specific trait transitions at 50 different mass classes. Stage are juveniles (J), yearlings (Y), non-reproductive adults (N), and reproductive adults (R).

### Supporting Material S4 - Population viability analyses

Fig. 4 in the main text shows results of population projections in which the averages and standard deviations of environmental quality, $Q_y$, were perturbed but kept fixed for the 50 years of projections. Fig. S4.1 shows results based on introducing a trend in the averages of $Q_y$, *i.e.*, a decrease by 0.01 each year of projections.

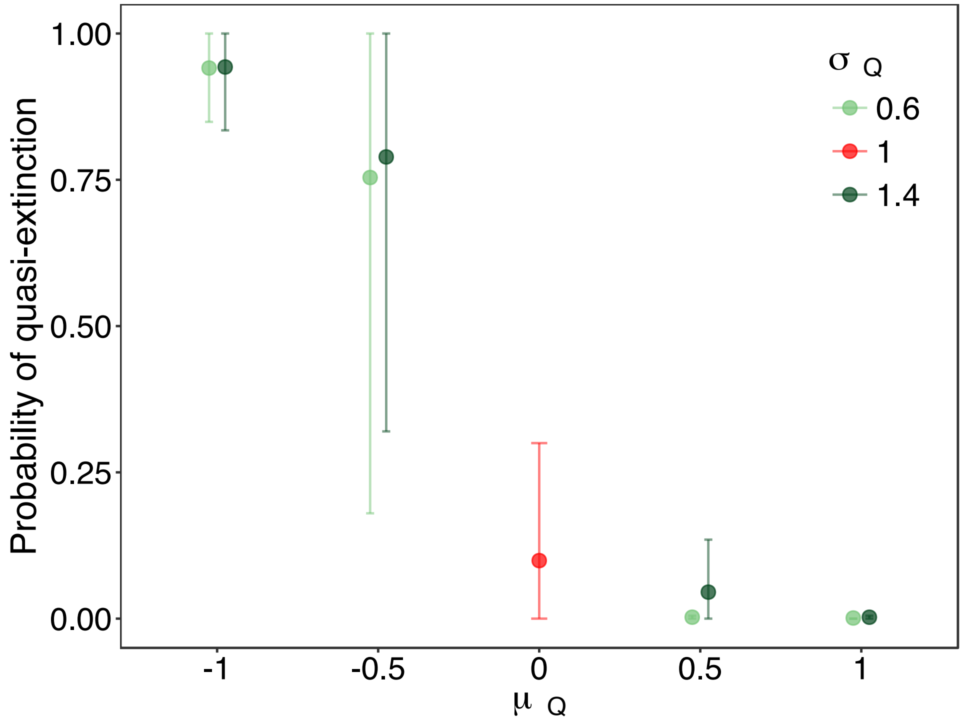


**Figure S4.1:** Probability of quasi-extinction (*i.e.*, < 4 non-juveniles in the population) of yellow-bellied marmots under different scenarios of environmental change. The scenarios consisted of projecting population dynamics for 50 years, each year decreasing the mean ($\mu$) of environmental quality $Q_y$ (by 0.01) in all demographic processes simultaneously; but fixing standard deviation ($\sigma$). Points and error bars show averages ± 95 % C.I. across 1,000 posterior parameter samples obtained from the Bayesian population model. Base simulations (fixing $\mu_Q = 0$; $\sigma_Q = 1$) are depicted in red.

To incorporate temporal autocorrelation into the simulations of population viability of the yellow-bellied marmot population, we defined a discrete Markov chain consisting of two states, (1) favorable or good environmental conditions and (2) unfavorable or bad conditions. The transitions between the two states were defined using the transition matrix:

$$\begin{bmatrix} p_g & p_b \\ 1 - p_g & 1 - p_b \end{bmatrix}$$

Here, $p_g$ and $p_b$ are the probabilities of transitioning to the good environment at time $t + 1$ when the environment was good or bad at $t$ respectively. These transitions can be derived from the long-term frequency of the good environment ($f$) and temporal autocorrelation ($AC$)(Tuljapurkar

& Haridas, 2006), where $p_b = f(1 - AC)$ and $p_g = AC + p_b$. In the simulations we used $f = 0.35$ and 0.65 and $AC$ = -0.5, -0.2, 0, 0.2, and 0.5.

Using all combinations of $f$ and $AC$ to define the Markov chain, we simulated environmental conditions for 50 years. We then linked these conditions to population dynamics by sampling values of the latent variable, $Q_y$, from a normal distribution with $\mu_Q = 1$ and $\mu_Q = -1$ when the environment was good and bad, respectively, and constructing winter and summer IPMs from these values. We kept the standard deviation, $\sigma_Q$, fixed at 0.6. We used two types of simulations of $Q_y$: fixing the $\mu_Q$ defining good and bad conditions for the 50 years of simulations and introducing a trend by decreasing $\mu_Q$ by 0.01 each year. Results of the simulations are shown in Fig. S4.2.

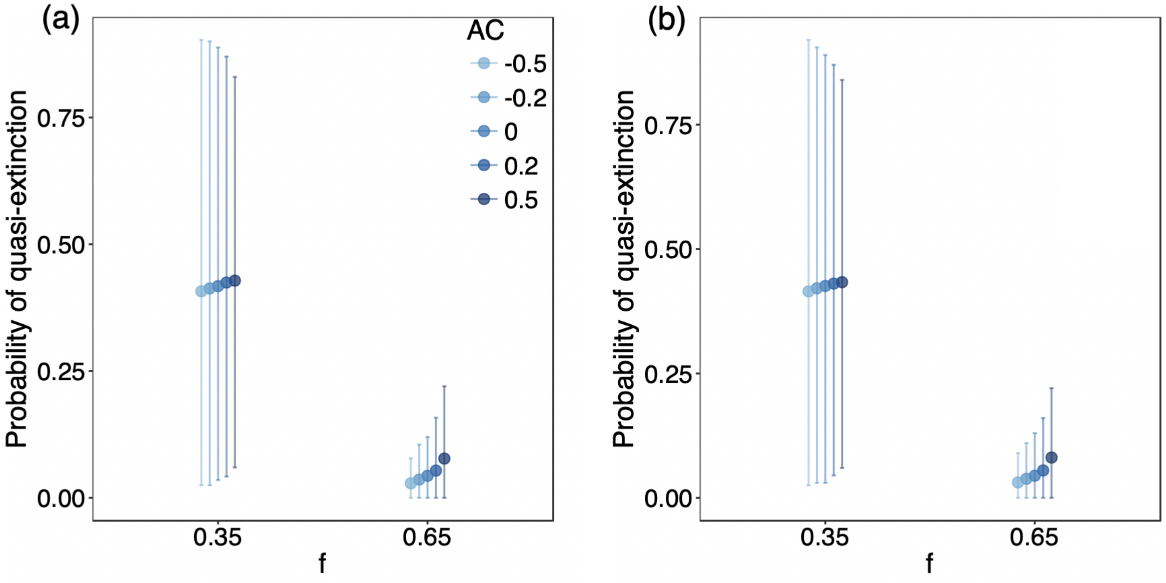


**Figure S4.2:** Probability of quasi-extinction (*i.e.*, < 4 non-juveniles in the population) of yellow-bellied marmots under autocorrelated environmental variation. Population dynamics were projected for 50 years, using two frequencies, $f$, of the good environmental state ($\mu_Q = 1$; $\sigma_Q = 0.6$) and different autocorrelation coefficients (AC). Each year, $\mu_Q$ either remained fixed (a) or decreased by 0.01 (b) in all demographic processes simultaneously. Points and error bars show averages ± 95 % C.I. across 1,000 posterior parameter samples obtained from the Bayesian population model.

### References

Caswell, H. (2001). *Matrix Population Models*. Sunderland: Sinauer.

Ellner, S. P., Childs, D. Z., & Rees, M. (2016). *Data-driven Modelling of Structured Populations: A Practical Guide to the Integral Projection Model*. Springer.

Griffith, A. B. (2017). Perturbation approaches for integral projection models. *Oikos*, 126, 1675–1686.

Haridas, C. V., & Tuljapurkar, S. (2005). Elasticities in variable environments: properties and implications. *The American Naturalist*, 166, 481–495.

Hindle, B. J., Rees, M., Sheppard, A. W., Quintana-Ascencio, P. F., Menges, E. S., & Childs, D. Z. (2018). Exploring population responses to environmental change when there is never enough data: a factor analytic approach. *Methods in Ecology and Evolution*, 9, 2283-2293.

Kaufman, C. G. & Sain, S. R. (2010). Bayesian functional ANOVA modeling using Gaussian process prior distributions. *Bayesian Analysis*, 5, 123–150.

Keller (2015). jagsUI: A Wrapper around 'rjags' to streamline 'JAGS' analyses. R package version 1.3. Available at: https://github.com/kenkellner/jagsUI. Last accessed November 25, 2019.

Kruschke, J. K. (2010). Bayesian data analysis. *Wiley Interdisciplinary Reviews. Cognitive Science*, 1, 658–676.

Legendre, P. & Legendre, L. (2012). *Numerical Ecology*, 3rd ed. Elsevier, London.

Paniw, M., Quintana-Ascencio, P. F., Ojeda, F., & Salguero-Gómez, R. (2017). Interacting livestock and fire may both threaten and increase viability of a fire-adapted Mediterranean carnivorous plant. *The Journal of Applied Ecology*, 54, 1884–1894.

Rees, M., & Ellner, S. P. (2009). Integral projection models for populations in temporally varying environments. *Ecological Monographs*, 79, 575–594.

Revelle, W. (2018). *psych: Procedures for Personality and Psychological Research*. Northwestern University, Evanston, Illinois, USA. Available at https://cran.r-project.org/web/packages/psych/index.html. Version 1.8.12. Last accessed November 25, 2019.

Schwartz, O. A., & Armitage, K. B. (2002). Correlations between weather factors and life-history traits of yellow-bellied marmots. *Proceedings of the 3rd International Marmot Conference, Cheboksary, Russia*.

Schwartz, O. A., & Armitage, K. B. (2005). Weather influences on demography of the yellow-bellied marmot (*Marmota flaviventris*). *Journal of Zoology*, 265, 73–79.

Tuljapurkar, S., & Haridas, C. V. (2006). Temporal autocorrelation and stochastic population growth. *Ecology Letters*, 9, 327–337.

Wood, S.N. (2006). *Generalized Additive Models: An Introduction with R*. Chapman and Hall/CRC, London.
